# Cytoglobin-dependent NO-sGC-cGMP signaling regulates ventricular morphogenesis and diastolic function

**DOI:** 10.64898/2026.03.13.711730

**Authors:** Adam Austin Clark, Rasmus Hejlesen, Tzu-Ting Weng, Muddassar Iqbal, Akueba Bruce, Paola Corti

## Abstract

**Aims:** Hypoplastic left heart syndrome (HLHS) is a severe congenital heart disease characterized by ventricular hypoplasia and impaired cardiac function. Clinically, inhaled nitric oxide (NO) therapy is used to reduce pulmonary vascular resistance and improve cardiopulmonary stability in HLHS patients. However, whether NO signaling contributes to HLHS pathogenesis remains unknown. Cytoglobin (Cygb) is a heme-protein traditionally thought to limit NO bioavailability and downstream activation of the soluble guanylate cyclase (sGC) cyclic guanosine monophosphate (cGMP) signaling pathway. Unexpectedly, our recent work shows that Cygb enhances NO signaling through activation of NO synthase, leading to downstream activation of sGC-cGMP signaling. In zebrafish embryos, Cygb-dependent NO signaling is required for normal cilia motility and the establishment of correct cardiac laterality. Here, our aim was to determine whether Cygb-dependent NO-sGC signaling linked to cilia function regulates cardiac morphogenesis and contributes to ventricular hypoplasia in HLHS.

**Methods and Results:** We found that loss of *Cygb* (*cygb2*) in zebrafish disrupts NO-sGC signaling during cardiogenesis, altering cardiac progenitor organization and migration within the anterior lateral plate mesoderm. Disruption of these processes impairs heart tube morphogenesis, thereby producing a compact ventricle wall characterized by increased wall thickness (despite preserved cardiomyocyte number) reduced ventricle size and decreased stroke volume, recapitulating key features of HLHS. Genetic disruption of the sGC α-subunit (*gucy1a1)* and pharmacological NO scavenging phenocopy the *cygb2* mutant phenotype, resulting in reduced cGMP levels, compact ventricular architecture and decreased stroke volume. Consistently, restoration of NO-sGC signaling in *cygb2* mutants rescues early cardiac progenitor patterning, ventricular morphology and stroke volume.

**Conclusions:** These findings identify Cygb-dependent NO-sGC signaling as a critical developmental pathway for ventricular development and performance, temporally linking cardiac progenitor dynamics to cilia-dependent signaling associated with left-right patterning. This study further suggests that pharmacological activation of sGC may provide a therapeutic strategy for hypoplastic ventricular disease.

**Graphical Abstract:** 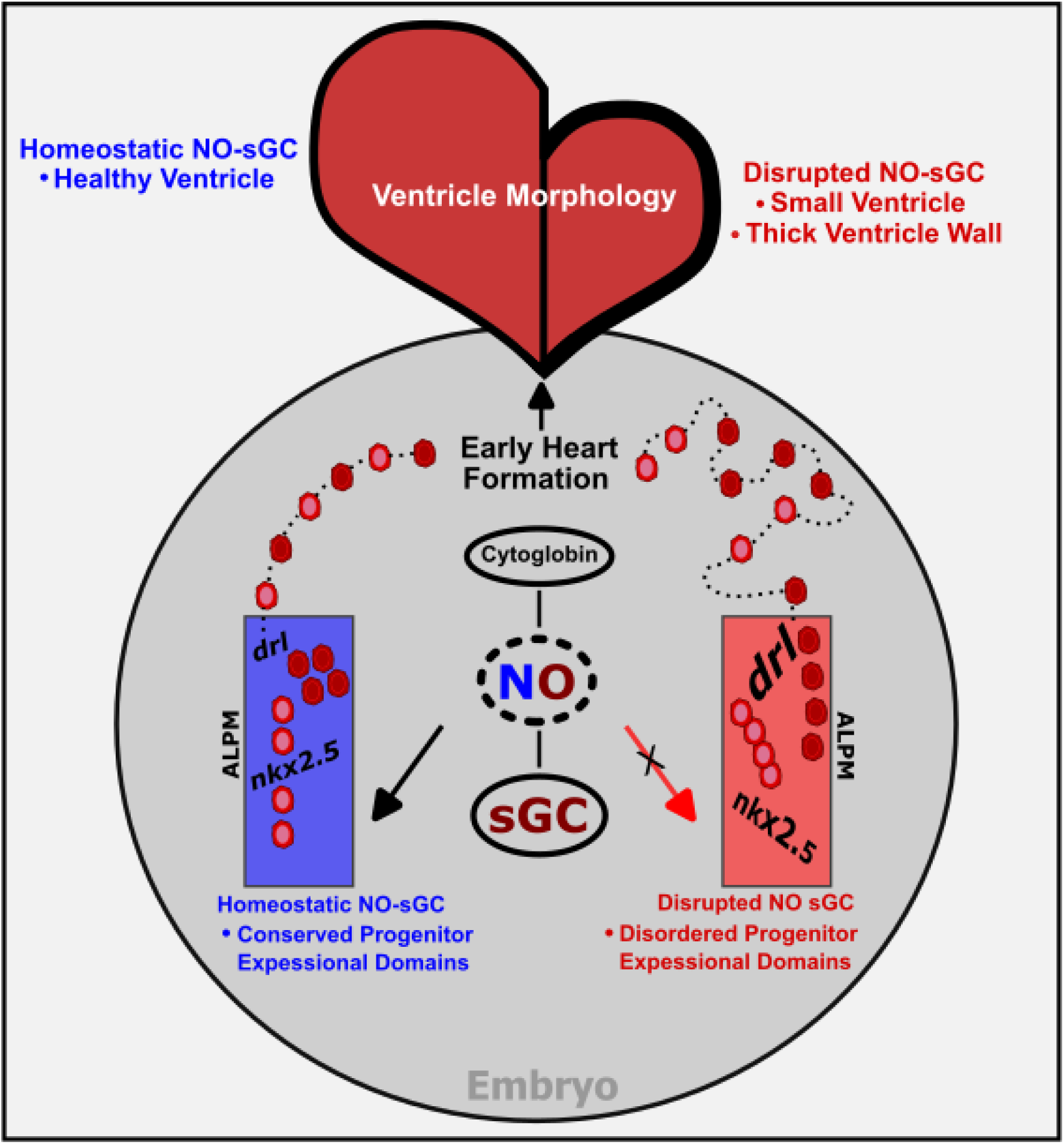

## 1. Introduction

Hypoplastic left heart syndrome (HLHS) is a complex congenital heart disease with a multigenic etiology.^1,2^ Familial clustering supports a genetic basis, yet the majority of causative genes remain unidentified. The phenotype is heterogeneous, variably affecting the left ventricle, valves, and aortic outflow tract, and no single-gene mutation has been shown to reproduce the full HLHS spectrum in animal models. Traditionally, HLHS was attributed to diminished intracardiac flow during development, leading to secondary hypoplasia. However, recent work has shifted this paradigm, suggesting that HLHS may arise from primary defects in myocardial and endocardial development, resembling a developmental cardiomyopathy in origin rather than a purely hemodynamic lesion.^3^ Additionally, an emerging body of evidence links HLHS to disturbances of left–right patterning and cilia function, suggesting a new developmental mechanism contributing to disease pathogenesis.^4^ Based on case reports and large registry-based analyses,^4,5^ HLHS has been observed in association with heterotaxy, situs inversus, gut malrotation, biliary atresia and splenic anomalies.^6–8^ These observations further support the concept that defects in laterality determination and ventricular morphogenesis may share developmental origins. According to recent interpretations, HLHS may originate from early intrinsic disruptions in cardiac progenitors arising during and prior to heart tube formation,^9^ and may occur in parallel with, or as a consequence of, abnormal left-right signaling. Findings from recessive forward genetic screens in fetal mice further support this view, revealing that cilia and cilia-transduced signaling pathways play central roles in HLHS pathogenesis.^10^

Integrative genetic analysis combining human sequencing with developmental datasets and mouse models of HLHS, identified CYGB as a candidate gene within a developmental gene network associated with the disease,^11^ suggesting a potential role for CYGB in pathways regulating early cardiac morphogenesis. CYGB is a hexa-coordinate heme-containing globin expressed ubiquitously in vertebrate tissues.^12,13^ Unlike its penta-coordinate relatives, hemoglobin and myoglobin, which primarily bind and transport oxygen, CYGB is more prone to function as a redox-active enzyme regulating NO metabolism through reversible redox cycling of its heme iron.^14^ The reduced heme iron can catalyze nitrite reduction to bioactive NO,^13,15^ while the oxygenated heme iron can dioxygenate NO to inert nitrate,^16–18^ thus maintaining oxygen-dependent NO homeostasis. While these reactions are possible in vitro, mouse studies indicate that the predominant physiological role of CYGB is to scavenge NO, modulating vascular tone through NO deoxygenation^19^ and to protect cells against oxidative stress via superoxide dismutase-like activity.^20^ Unexpectedly, we discovered a new critical role for Cygb in development, where Cygb instead enhances NO signaling. The zebrafish *Cygb* (*cygb2)* knockout exhibits cardiac laterality defects due to impaired NO signaling, a phenotype characteristic of human disorders associated with defective cilia function. Mechanistically, we demonstrated that Cygb2 localizes to cilia within the zebrafish laterality organ (Kupffer’s vesicle, KV), and increases NO bioavailability through activation of the nitric oxide synthase enzyme (Nos2b).^21^ Loss of *cygb2* disrupts ciliary structure, motility and directional fluid flow, resulting in impaired KV activity and cardiac laterality defects.^21^ Whether this function is directly linked to cardiac morphogenesis mimicking HLHS remains unknown and the extent to which Cygb-dependent regulation of laterality intersects, potentially through ciliary mechanisms, has not been explored.

Zebrafish cardiac development initiates within the anterior lateral plate mesoderm (ALPM), where bilateral populations of mesodermal progenitors undergo progressive spatial organization prior to overt myocardial differentiation.^22,23^ Between the 14–16 somite stages (ss), these progenitors undergo tightly coordinated mediolateral convergence and anterior–posterior elongation to assemble a coherent cardiac disc that subsequently develops into the linear heart tube.^23,24^ At these stages of cardiac development, ventricular formation is exquisitely sensitive to perturbations in progenitor number and spatial positioning as even subtle defects in coordinated cell migration or tissue architecture can give rise to abnormalities in ventricular morphogenesis, whereas atrial development may remain relatively preserved due to its later progenitor contribution.^24,25^ Importantly, these morphogenetic events critical to ventricular development occur immediately downstream of KV mediated left–right axis establishment,^23,26^ raising the possibility that laterality patterning cues coordinating embryonic asymmetry also influence spatial patterning and migratory behavior of cardiac progenitors, shaping ventricular growth and chamber morphogenesis. One of the earliest markers of cardiac progenitors in zebrafish is *draculin* (*drl*), a transcription factor activated during gastrulation that delineates bilateral cardiac progenitor fields within the ALPM.^23,27^ Alterations in ALPM spatial geometry, such as anterior– posterior elongation of the *drl* domain, reflect early disruption of progenitor distribution or specification,^23^ defects expected to impair migratory cues and ventricle assembly. As development proceeds, myocardial *drl* expression is restricted to first heart field-derived cardiomyocytes, providing a readout of cardiac disc formation and heart tube assembly, stages that directly precede ventricular chamber growth and functional maturation.^23^

*NKX2 homeobox 5* (*Nkx2.5)* encodes a conserved cardiac homeobox transcription factor expressed within the forming cardiac disc at the transition from migratory cardiac progenitors to differentiated myocardium.^23,27^ In both mouse models and human patients, dysregulation of NKX2.5 activity is associated with congenital heart diseases and hypoplastic ventricular phenotypes^28^. In the zebrafish embryo, *nkx2.5* expression is conserved and is initially induced in bilateral ALPM domains as cardiac progenitors converge toward the midline, coinciding with progenitor stabilization and the initiation of chamber-specific transcriptional programs.^29^ By 28 hpf, *nkx2.5* expression marks specified cardiomyocytes within the heart tube, predominantly ventricular myocardium,^24^ such that its expression at this stage reflects proper myocardial differentiation and maintenance of cardiac identity.

While Cygb-dependent regulation of embryonic laterality is now established,^21^ its role in coordinating cardiac progenitor migration and myocardial differentiation remains unknown. Here, we demonstrate that *cygb2* deficiency in zebrafish disrupts early cardiac progenitor organization and ventricular chamber formation, producing a small, compact ventricle with reduced diastolic filling. These features parallel ventricular hypoplasia and impaired diastolic function observed in HLHS. Crucially, these defects are phenocopied in sGC mutants and fully rescued by pharmacological activation of NO-sGC signaling, identifying a developmental pathway that links ciliary NO regulation to cardiac progenitor morphogenesis and ventricular growth.

## 2. Methods

All animal studies were approved by the University of Maryland Institutional Animal Care and Use Committee (IACUC-1022015), Baltimore, MD, USA. The programs and facilities are USDA registered and covered under an assurance with the Office of Lab Animal Welfare (OLAW). Procedures followed guidelines outlined by the NIH Guide for the Care and Use of Laboratory Animals (8th Edition, 2011). Embryos were maintained at 28.5 °C until the desired developmental stage. Animals were euthanized according to approved protocols; embryos >3 days post fertilization (dpf) to adult (anesthetic overdose 300mg/L buffered tricane), embryos <3dpf (paraformaldehyde immersion).

### 2.1 Animal Studies

For all experiments, clutch-matched embryos from dome stage (∼ 4hpf) to 5dpf were randomly assigned to treatment groups; sex cannot be determined at these stages. The AB (WT) genetic background was used for generation of mutant and transgenic lines. Transgenic lines in this study include Tg(*myl7:EGFP*) and Tg(*tbx5a:EGFP*; *tcf21:dsRED*). The *cygb2^801a^* and the *gucy1a1^9bp^* mutants were generated using CRISPR-Cas9 genome editing as previously described^21^ and guide RNAs are listed in^21^ and *Table S1*. Embryos were raised and screened for restriction enzyme site disruption (N1aIII) using primers specific to exon 4 of *gucy1a1* (*Table S1*). The Tg(*myl7:EGFP*) and Tg*(tbx5a:EGFP);*Tg*(tcf21:dsRED)* lines were crossed with *cygb2^801a^* to produce *cygb2^801a^*;Tg*(myl7:EGFP)*, *cygb2^801a^*;Tg*(tbx5a:EGFP);*Tg*(tcf21:dsRED),* and wildtype sibling lines.

### 2.2 Stroke Volume

Stroke volumes were calculated from live imaging. To calculate end-diastolic volume (EDV) and end-systolic volume (ESV) frames were identified from high-speed wide-field (bright-field and fluorescent) time-lapse videos of beating zebrafish hearts at the moments of maximal and minimal ventricular cavity size. The long-axis length (*L*) and short-axis diameter (*D*) of the ventricle were measured from frames using FIJI imaging software. EDV and ESV were calculated using the established prolate spheroid model^30^ according to the formula:

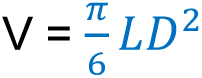

L represents ventricular long-axis length and D the diameter at mid-ventricle (*Figure S1*). Stroke volume (SV) was derived as SV = EDV − ESV. Ventricle wall thickness was calculated at maximal and minimal cavity size, identified from high-speed differential interference contrast (DIC) microscopy videos of beating zebrafish hearts. The measurable area and length of the ventral-facing ventricle wall was calculated using FIJI imaging software and the polygon selections and segmented line tool. The area of the selected polygon was divided by the length of the measured wall to generate a ratio-based metric of thickness. Bright field and fluorescent images were captured using a Zeiss Axio Observer inverted microscope with attached Axiocam 807 mono camera and a Leica M165FC with attached K3C camera.

### 2.3 Ventricle Volume and morphology

Whole mount confocal z-series and compiled z-projections were collected on paraformaldehyde fixed zebrafish embryos from 16ss to 3 days post fertilization. Immuno-labeled embryos were mounted on glass slides and imaged on a Nikon Ti2 inverted W1 spinning disk confocal microscope with attached Hamamatsu sCMOS camera. Collected z-series were compiled into max projections and 3D objects with surface renderings using FIJI and Imaris imaging software. Ventricle volume and wall thickness were calculated using thresholding on selected regions of interest (ROI). Z-slices covering full depth and width of ventricle were compiled, and images were converted to 8-bit and set to grey scale. To calculate the volume of myocardium, thresholding was set to black and white, area measurements for signal-positive region were measured for each z-slice, summed, and multiplied by the distance between z-slices. Ventricular lumen volume was calculated by measuring the area of signal-negative region within the surrounding signal-positive region, summing the areas across z-slices and multiplying by z-slice distance. To create a ratio-based metric to measure thickness across a 3-dimensional structure, myocardium volume was divided by lumen volume. Cell density was calculated by dividing the total cardiomyocyte number by the myocardium volume. Total ventricle cardiomyocyte number was calculated using the collected z-series and FIJI imaging software. Z-slices spanning the full depth and width of the ventricle were compiled, and analyzed frame by frame, and EGFP-positive cardiomyocytes were marked, tracked, and counted. Ventricle volume at 5 dpf was generated from z-series collected using a Zeiss LS7 Lighsheet confocal microscope. Ventricle area of z-slices was calculated by thresholding (Huang Over Under) of the EGFP positive region within a ROI defining the ventricle including the luminal space. Using FIJI and open-source macro code ventricular volume was calculated by multiplying the summed areas by the distance between z-slices. Progenitor cell fusion, cardiac disc, and heart tube areas were calculated using thresholding of EGFP and CY5 positive areas in selected ROIs of max z-projections collected using spinning disk confocal microscopy.

### 2.4 In situ hybridization

Whole-mount in situ hybridization was performed on paraformaldehyde fixed zebrafish embryos as previously described^21^ at 10-12ss and 28 hpf using riboprobes targeting *drl*, *kinase insert domain receptor like* (*kdrl*), and *nkx2.5*. Transcripts were amplified by PCR and cDNA was obtained from pooled embryos using transcript specific primers (*Table S1*). Images were collected using bright-field microscopy on a Zeiss Discovery V.8 SteReo stereoscope with attached Axiocam 503 color camera, brightness and contrast were unaltered, and images were formatted using FIJI imaging software. To determine expression patterning variance, raw counts of phenotype rankings were used for statistical comparisons with the Fishers exact test, percentages were plotted for graphical representation, ns - not significant ≥ 0.05, * P≤ 0.05, ** P≤ 0.01, *** P≤ 0.001, **** P≤ 0.0001.

#### Drug Exposure

Carboxy-PTIO potassium salt (2-4-carboxyphenyl-4,4,5,5-tetramethylimidazoline-1-oxyl-3-oxide; cPTIO, Sigma–C221) and BAY 582667 (Cinaciguat, Bay58, Sigma–SML1532) were diluted in DMSO and used at final concentrations of 500µM and 120µM respectively. (Z)-1-(N,N-diethylamino)diazen-1-ium-1,2-diolate (DETA NONOate) (Cayman-82100) and (Z)-1-[N-(2-aminoethyl)-N-(2-ammonioethyl)amino]diazen-1-ium-1,2-diolate (PAPA NONOate) (Cayman-2140) were resuspended in sodium hydroxide and diluted in phosphate buffered saline at a final concentration of 250µM and 5.73µM. Drug exposure was achieved by incubation in embryo medium (E3).

#### Immunohistochemistry

Whole-mount immunohistochemistry was performed as previously described^21^ using primary antibodies for myosin heavy chain 6 (Developmental Studies Hybridoma Bank S46). Embryos were mounted on glass slides with Vectashield Hardset Antifade Mounting Medium (H-1500-10) and imaged on a Nikon Ti2 inverted W1 spinning disk confocal microscope equipped with an Hamamatsu sCMOS camera.

#### cGMP ELISA

cGMP levels in zebrafish embryos were measured by competitive ELISA following manufactures protocol (Enzo-ADI-900-164). Embryos at 28 hpf were incubated in 250µM DETA NONOate for 30 minutes at 28.5 °C. Fifty embryos per sample were pooled and homogenized in 0.1M hydrochloric acid on ice. Lysates were centrifuged and the supernatant was collected for analysis. Samples were assayed in triplicate at two dilutions. cGMP concentration was calculated based on interpolation to the standard curve using a 95% confidence interval.

#### Cell culture

The human type II alveolar epithelial cell line A549 was kindly provided by Dr. Alan Cross (Professor, Department of Microbiology and Immunology, University of Maryland School of Medicine). A549 cells were cultured in RPMI-1640 media (Gibco-11875093) containing Penicillin/Streptomycin (Sigma-P4333) and fungizone (Amphoterecin B, Gibco-15290-026), and 10% fetal bovine serum (FBS) (Sigma-F4135) at 37^0^ Celsius and 5% carbon dioxide. Cell cultures were passaged a minimum of three times before performing experiments.

#### Small interfering RNA

Dicer Substrate RNA duplexed small interfering RNAs si-CYGB 13.1 and si-CYGB 13.2 targeting human *CYGB* were selected using Integrated DNA Technologies “Predesigned DsiRNA Selection Tool”. DsiRNA-mediated gene silencing was validated through transfection of sub-confluent A549 monolayers using Lipofectamine RNAiMAX transfection reagent (Thermo-13778030), non-targeting and human *CYGB* targeting siRNAs for 48 hours. Si-CYGB 13.1 showed higher protein knockdown in Western Blot experiments and was chosen for migration assays. To assess cell viability, transfected cells were pelleted and resuspended in a 1:1 mixture of cell suspension and 0.4% Trypan Blue. Cells were counted using a hemocytometer in a 0.05µm section. Cells excluding Trypan Blue were counted as live and cells stained with Trypan Blue were counted as dead, viability was calculated as a percent of live cells to total cell number.

#### Western Blot

A549 cells were lysed on ice in RIPA buffer (Sigma-R0278) containing protease inhibitor (Halt 1X, Thermo-1860932). Lysates were centrifuged at 12,500 RPM for 20 min at 4°C. The BCA assay was used to determine total protein concentration. Samples were loaded for gel electrophoresis in 4-20% precast gels (Bio-Rad-4561094) using 100 µg total protein per sample. Following electrophoresis protein was transferred to polyvinylidene fluoride membranes. Membranes were blocked with 3% bovine serum albumin (BSA) for 2 hours at room temperature under gentle agitation. Membranes were probed with primary rabbit anti-CYGB (Sigma-HPA017757) diluted at 1:1000 in 1% BSA and 0.1% Triton X100 in phosphate buffer saline (PBST) overnight at 4^0^ Celsius. Following 3 consecutive 5-minute washes at room temperature in PBST membranes were probed with anti-Rabbit IgG-HRP (Abcam-131366) at 1:10,000 in PBST with 1% BSA for 1 hour at room temperature. Following 3 consecutive 5-minute washes at room temperature in a solution of PBST protein bands were visualized using Clarity Max Western ECL substrate (Bio-Rad-1705060) following manufacturer’s protocol and a Bio-Rad ChemiDoc MP imaging box. Following 3 consecutive 5-minute washes with PBST at room temperature membranes were stripped using Restore Western Blot Stripping Buffer (Thermo-21059). Blocking and probing procedure was repeated using primary rabbit anti-β-Actin antibody (Sigma-A2066). Band intensity was normalized to β-Actin and quantified using FIJI imaging software.

#### Wound healing assay

A549 cells were seeded in 24-well plates at a density of 170,000 cells per well. After 24 hours cells were transfected with 2.5pmol non-targeting DsiRNA, 1.25pmol and 2.5 pmol si-CYGB 13.1. At 48 hours post transfection, a scratch was introduced with a 10µl pipette tip. Following the scratch, cells were washed with sterile 1x PBS and supplemented with RPMI-1640 media containing 10% (FBS). Cultures were imaged at 0 and 24 hours post scratch on a Leica DM IL LED inverted microscope using a 4X objective. Wound healing was assessed as percent migration into the scratch area relative to the migration of control. Image analysis was conducted using FIJI imaging software.

### 2.5 Statistical analysis

Unless otherwise noted data are presented as mean ± SD, and statistical significance was determined using an unpaired two-tailed *t*-test (or Mann–Whitney test for non-normal distributions), ns - not significant ≥ 0.05, * P≤ 0.05, ** P≤ 0.01, *** P≤ 0.001, **** P≤ 0.0001.

## 3. Results

### Cytoglobin is required for ventricular development and cardiac function

We previously defined a role for Cygb2 in cardiac laterality via regulation of cilia function. Here, we investigated whether Cygb2 is required for cardiac morphogenesis and performance and whether these are linked to laterality establishment. To visualize cardiac morphology, we generated a *cygb2* mutant reporter line by crossing the previously characterized *cygb2^801a^* mutant^21^ with a transgenic reporter line expressing EGFP under the control of the *myosin light chain 7* (*myl7)* promoter.^23^ In zebrafish, *myl7* expression begins at approximately the 16ss stage in cardiomyocyte progenitors regardless of eventual chamber contribution to the atrium or ventricle.^23^ We analyzed *myl7:EGFP* expression at 3 days post fertilization (dpf), when cardiac morphology is established, and the ventricle and atrium exhibit coordinated contraction and robust cardiac function. We first quantified the area of both cardiac chambers and observed a significant decrease in the ventricle of *cygb2^801a^* embryos compared to wildtype controls, (*Figure 1A–C, Movies S1,2*), while atrial area remained unchanged (*Figure 1D*). To precisely assess performance, we measured end diastolic volume (EDV) and end systolic volume (ESV) from in vivo high-speed recordings and calculated stroke volumes (SV) using the prolate spheroid method *(Figure S1).* Ventricles of *cygb2^801a^* embryos showed a marked decrease in SV and EDV with no change in ESV (*Figure 1E–H*), consistent with reduced diastolic filling rather than systolic impairment. As *cygb2^801a^* embryos exhibited cardiac laterality defects, we tested whether this ventricular defect might arise secondarily from laterality disturbances. We compared embryos with normal (situs solitus) and reversed (situs inversus) cardiac orientation within each genotype and detected no differences in SV, *cygb2^801a^* embryos displayed reduced SV relative to wildtype independent of cardiac left-right orientation (*Figure S2A–F*). Assessment of SV at 5dpf revealed that this phenotype persisted beyond initial ventricle morphogenesis (*Figure S3A–D, Movies S3-4*). Additionally, ventricle volume was measured using light sheet confocal microscopy on fixed *myl7:EGFP* samples and further supported the reduced ventricle volume observation and persistence at 5 dpf in *cygb2^801a^* compared to wildtype (*Figure S3E*). Collectively, these findings indicate a stable structural hypoplasia rather than a transient developmental delay. Despite reduced chamber size, ventricular cardiomyocyte number was unchanged (*Figure 1 I-J*). Instead, cells occupied a smaller volume, yielding a more compact myocardium and a thicker ventricular wall (*Figure 1K–M*). These data indicate that *cygb2* deficiency primarily affects ventricular morphogenesis and chamber remodeling, not cardiomyocyte proliferation or survival. The reduced, thick-walled ventricle observed in *cygb2^801a^* embryos accompanied by reduced SV mirrors key structural features of HLHS thus *cygb2* deficiency may represent a mechanistically relevant model of HLHS-like ventricular hypoplasia.

**Figure 1.**
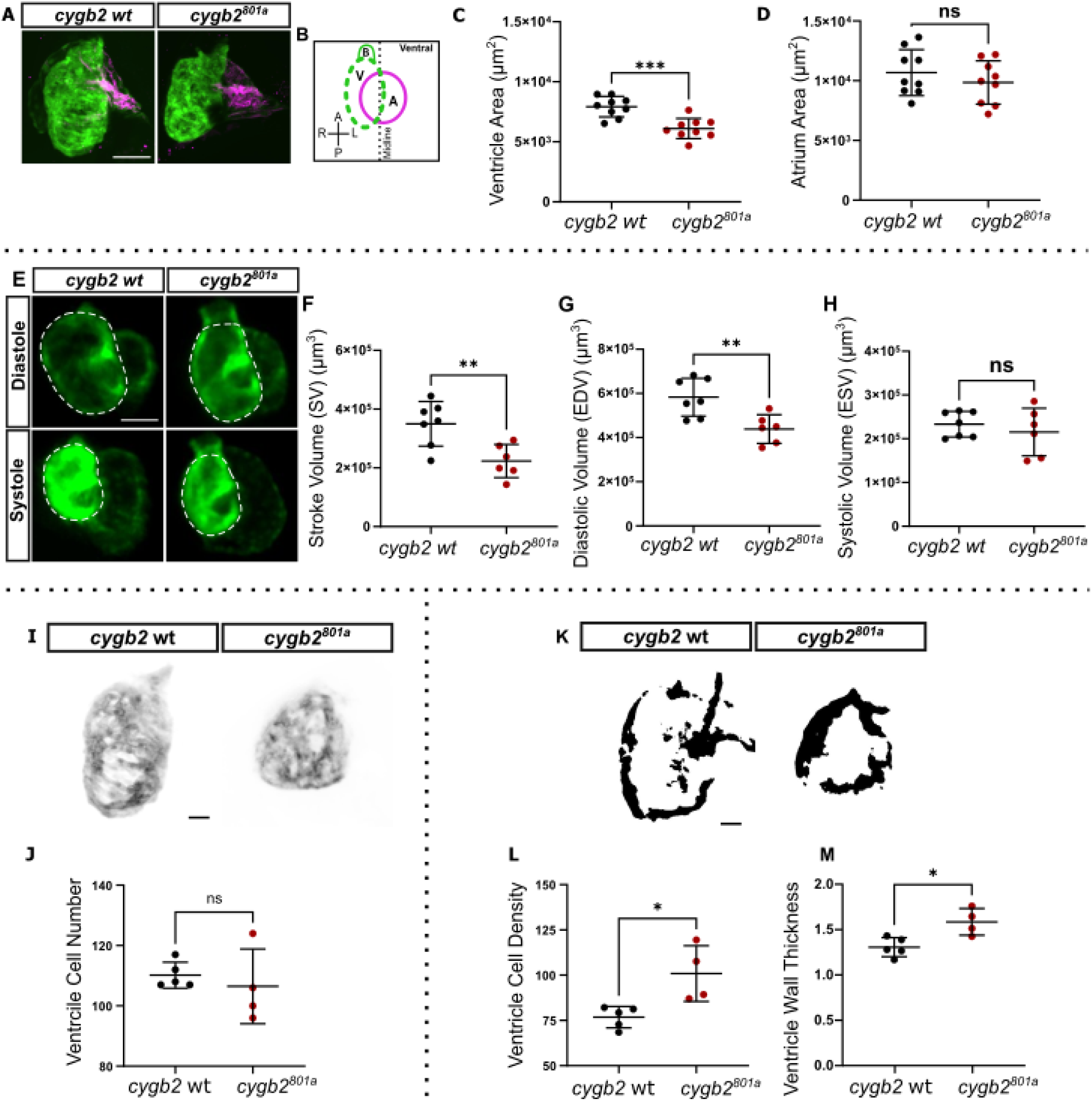
Loss of *cygb2* leads to ventricular hypoplasia and reduced cardiac output. **(A)** Representative confocal projections of *cygb2wt*;Tg*(myl7*:EGFP) and *cygb2^801a^*;Tg*(myl7*:EGFP) hearts at 3 days post fertilization (dpf), immuno-labeled for Myh6 (EGFP, green; Myh6, magenta). Scale bar, 50µm **(B)** Schematic illustrating heart chamber anatomy and cardiac orientation. A-atrium; V-ventricle; B-bulbus arteriosus **(C,D)** Comparison of ventricle and atrium area. **(E)** Representative frames from fluorescence time-lapse imaging of live *cygb2wt*;Tg*(myl7*:EGFP) and *cygb2^801a^*;Tg*(myl7*:EGFP) embryos 3 dpf showing ventricle volume at diastole and systole, dashed white line demarcates the ventricle (EGFP, green). Scale bar, 50 µm. **(F-H)** Comparison of stroke volume, diastolic volume and systolic volume. **(I)** Representative confocal projections of ventricles at 3 dpf, images are greyscale converted and inverted to highlight cardiomyocyte density. Scale bar, 20µm. **(J)** Comparison of ventricle cell number. **(K)** Representative confocal z-slices at ventricular midpoint. Images were converted to greyscale signal positive area (ventricular myocardium) shown in black. **(L)** Comparison of cardiomyocyte density. **(M)** Comparison of ventricular wall thickness. Sample size (n) corresponds to independent embryos represented by individual data points on plots. Student t-Test; *ns*, not significant; *p<0.05; **p<0.01;***p<0.001

### Nitric oxide restores ventricular size and function in *cygb2* mutants

We previously showed that Cygb2 regulates cardiac laterality determination^21^ through NO signaling. To test whether the reduced SV in *cygb2^801a^* embryos depends on NO, we first confirmed the acute physiological action of NO donors. Ten-minute exposure at 78 hpf to the pharmacological NO donor DETA-NONOate (DETA) increased SV in both wildtype and *cygb2^801a^* embryos *(Figure S4A,B),* consistent with the reported hemodynamic effect of NO in teleost fish larvae^31^ where the increase in SV results from reduced afterload due to relaxation of the outflow vasculature. A fast-releasing NO donor, PAPA-NONOate (PAPA), produced a similar acute increase, validating the assay (*Figure S4C,D*). To determine whether the altered ventricular developmental defects could be rescued by NO supplementation, we tested the effect of chronic DETA treatment. DETA was administered in a vehicle (sodium hydroxide), which alone did not affect SV (*Figure S4E*). We performed DETA treatments from the 2ss stage to 3 dpf. Chronic exposure produced opposite outcomes depending on genotype, in *cygb2^801a^* embryos ventricular size increased with consequent rescue in SV, whereas in wildtype embryos ventricular size was reduced, resulting in decreased SV (*Figure 2A,B*). Because prolonged NO exposure affects cardiac morphogenesis rather than acute mechanics, these bidirectional responses under identical dosing indicate that *cygb2* mutants possess a specific deficit in NO-dependent developmental signaling, while excess NO in wildtype embryos disrupts the finely tuned program governing ventricle morphogenesis. To provide further evidence supporting NO dependence, we treated embryos at 2ss with the NO scavenger cPTIO to phenocopy the cardiac phenotype observed in *cygb2* mutants (*Figure 2A,C*). As expected, ventricular volume was reduced by NO scavenging in wildtype embryos and was further exacerbated in *cygb2^801a^* embryos, consistent with a requirement for NO signaling in normal ventricular development. A time course initiating DETA at stages from dome to 12ss showed maximal rescue when treatment began at dome stage to 2ss, diminishing by 12ss (*Figure 2D*), suggesting Cygb2-dependent NO signaling becomes critical for cardiac specification and morphogenesis during early somitogenesis. Together, these findings demonstrate that *cygb2* is required to maintain appropriate NO levels during early cardiac progenitor organization, thereby ensuring proper ventricular morphogenesis and SV.

**Figure 2.**
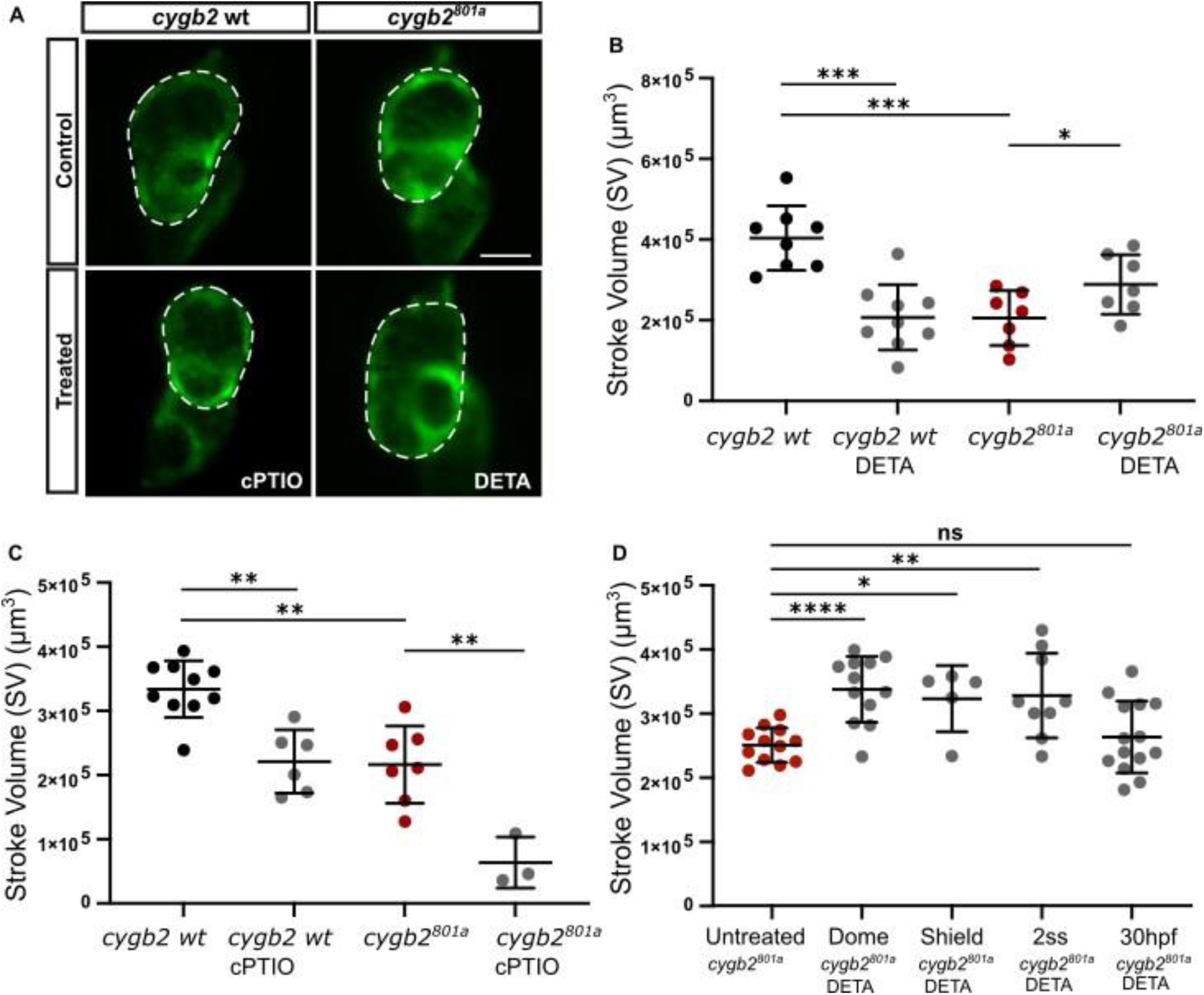
NO rescues the *cygb2* mutant phenotype, whereas NO scavenging phenocopies the mutant. **(A)** Representative frames from fluorescent time-lapse imaging of live *cygb2wt*;Tg(*myl7*:EGFP) and *cygb2^801a^*;Tg(*myl7*:EGFP) mutants at 3 days post fertilization (dpf), showing the effect of treatment with a pharmacological NO scavenger (cPTIO) and NO donor (DETA) on diastolic volume (EGFP, green). Scale bar, 50µm. **(B)** Comparison of stroke volume at 3dpf in untreated control and NO donor (DETA) treated, and **(C)** NO scavenger (cPTIO) treated embryos, treatments starting at 2 somite stage (ss). **(D)** Comparison of stroke volumes following chronic treatments starting at different developmental stages (dome, shield, 2ss, and 30hpf). Sample size (n) corresponds to independent embryos represented by individual data points on plots. Student t-Test; *ns*, not significant; *p<0.05; **p<0.01;***p<0.001; ****p<0.0001.

### Loss of Cygb2 disrupts early cardiomyocyte migration

Given that the ventricular phenotype observed in *cygb2^801a^* embryos originates during early development, we next examined cardiac morphogenesis to determine when these defects first emerge. In zebrafish, cardiac progenitors originate bilaterally within the ALPM and progressively migrate toward the midline, where they fuse to form the cardiac cone and subsequently the primitive heart tube. To assess whether this process was altered in *cygb2* mutants, we analyzed the distribution of *myl7*-positive cardiomyocyte precursors during heart field assembly. At 16ss when cardiomyocyte progenitors have initiated migration but have not yet fused, *cygb2^801a^* embryos displayed a significantly broader *myl7*-positive domain compared with wildtype, indicating an expanded or delayed coalescence of cardiac progenitors at this stage (*Figure 3A-C*). By 20ss, when the cardiac disc becomes morphologically visible at the midline, the *myl7*-positive region remained wider and less compacted in *cygb2^801a^* embryos, occupying a larger total area relative to controls (*Figure 3D-F*). These results suggest that Cygb*2* is required for the coordinated medial convergence of cardiomyocyte progenitors during heart field assembly. As development progressed, the cardiac tube of *cygb2^801a^* embryos remained visibly smaller than in wildtype. At 34 and 48 hpf, the ventricular area was significantly reduced indicating that the morphogenetic defect persists through the stages of heart tube elongation and chamber ballooning, whereas the atrium appeared morphologically normal throughout (*Figure S5A-H).* This suggests a ventricle-specific effect consistent with the smaller, compact ventricular morphology observed at later stages. To establish NO as a critical regulator throughout ventricular morphogenesis, embryos were treated with the NO donor DETA beginning at 2ss. These treatments restored *myl7*-positive cardiac field at 16 and 20ss (*Figure 3A-F)* and rescued ventricular area at 34 and 48 hpf (*Figure S5A-H*) to near wildtype levels. These findings demonstrate that *cygb2* is essential for the proper migration and coalescence of cardiac progenitors and for subsequent expansion of the ventricular chamber, and that both processes depend on NO-mediated signaling.

**Fig. 3.**
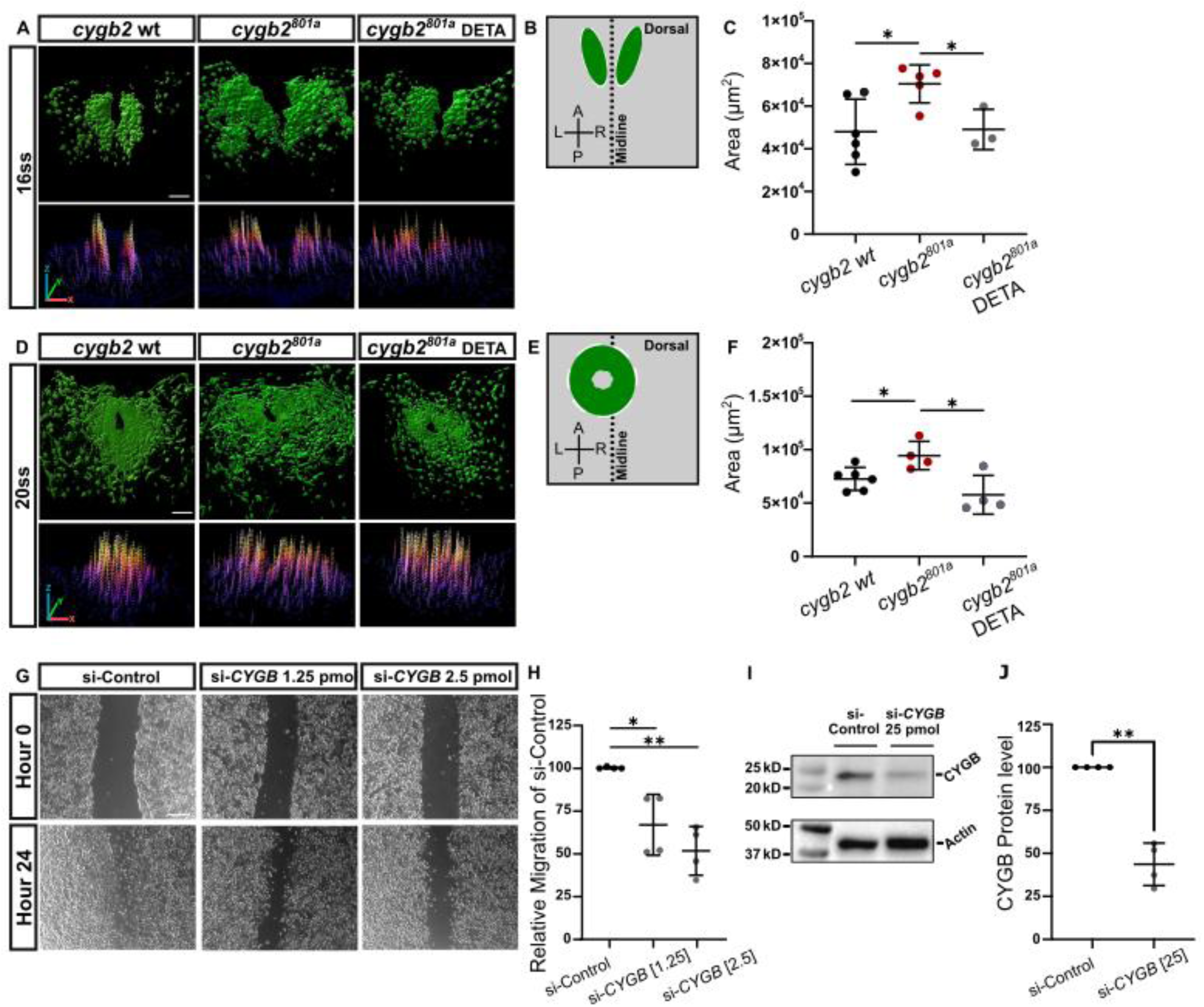
*cygb2* mutants show defective cardiac progenitors migration, and CYGB knockdown inhibits cell migration in human airway epithelial cell culture. **(A,D)** 3D surface rendering and surface plots from *cygb2wt*;Tg(*myl7*:EGFP) and *cygb2^801a^*;Tg(*myl7*:EGFP) showing cardiac progenitors at midline convergence, 16 somite stage (ss), and cardiac disc, 20 ss, renderings and surface plots generated from whole-mount confocal z-series (EGFP, green). NO donor (DETA) treatment starting at 2ss. Scale bar, 50µm. (**B,E**) Schematics illustrating cardiac progenitor organization and embryonic orientation. **(C,F)** Effect of chronic DETA treatment on cardiac progenitor organization, represented by EGFP positive area. Treatment starting at 2ss. **(G)** Representative bright-field images of human airway epithelial A549 cultures transfected with a non-targeting (si-Control) and *CYGB* targeting (si-CYGB 13.1) siRNAs at 0 and 24 hours post scratch. **(H)** Comparison of percent migration between si-Control and si-*CYGB* at 1.25 pmol and 2.5 pmol. **(I)** Representative images of Western blot bands for CYGB and β-Actin. **(J)** Percent CYGB protein knockdown with si-*CYGB* at 25 pmol relative to si-Control. **(A-F)** Sample size (n) corresponds to independent embryos **(G-J)** Sample size (n) corresponds to independent cultures, all represented by individual data points on plots. Student t-Test; *p<0.05; **p<0.01.

Upon observing aberrant organization of cardiac progenitors during midline convergence and cardiac disc formation in *cygb2^801a^* embryos, we directly tested whether cell migratory behavior was disrupted using an in vitro wound-healing assay in the human type II alveolar epithelial A549 cell line. *CYGB* transcript knockdown by small interfering RNA caused a dose-dependent reduction in cell migration into the wound area (*Figure 3G,H*). CYGB protein knockdown efficiency and cell viability were confirmed by Western Blot analysis (*Figure 3I,J and S6A*) and viability assay (*Figure S6B*). These data suggest that abnormal cardiac progenitor contribution to early heart structures may result from disruption of cellular machinery regulating cell migration and/or cell adhesion and further indicate that Cygb-NO signaling influences migratory behavior across cell types and species.

### Soluble guanylate cyclase (sGC) is required for ventricle development and heart function

NO can signal through multiple mechanisms, including protein S-nitrosylation, nitration reactions and formation of reactive nitrogen species. However, our data indicate that here NO primarily signals through its canonical receptor sGC. Upon ligand binding, NO-sGC catalyzes the conversion of guanosine triphosphate (GTP) to cGMP, thereby initiating secondary downstream signaling events^32^. To test whether the cardiac phenotype observed in *cygb2^801a^* embryos depends on sGC, we generated an sGC mutant (*gucy1a1^9bp^*) using CRISPR/Cas9-mediated site-directed mutagenesis. Genomic DNA from embryos derived from a maternal zygotic mutant line revealed a 9 bp deletion in exon 4 of the sGCα1 subunit, which was confirmed at the genomic and mRNA level by Sanger Sequencing (*Figure S7A*). Loss of the normal transcript was validated by in situ hybridization using a 300 bp probe spanning the edited region, which showed markedly reduced hybridization signal in *gucy1a1^9bp^* embryos compared to wildtype, whereas a 1000 bp probe located downstream of the edited site produced comparable staining in both mutants and wild type (*Figure S7B*). Functional loss of sGC activity in *gucy1a1^9bp^* embryos was confirmed by failure of DETA to restore cGMP levels in *gucy1a1^9bp^* compared to wildtype (*Figure S7E*). In contrast, DETA treatment restored cGMP levels in *cygb2^801a^* embryos,^21^ supporting the interpretation that Cygb2 functions upstream of sGC to regulate NO-dependent cGMP production during cardiac development. To determine whether sGC disruption phenocopies the *cygb2^801a^* cardiac defects observed here, we quantified ventricular performance in *gucy1a1^9bp^* embryos using DIC microscopy and high-speed imaging. SV was significantly reduced in *gucy1a1^9bp^* compared with wildtype embryos, driven by a decrease in EDV while ESV remained unchanged (*Figure 4A-E, Movies S5,6*). This pattern mirrors the functional phenotype of *cygb2^801a^* embryos, indicating that loss of sGCα1 function recapitulates the diastolic filling defect observed in the *cygb2* mutant. Furthermore, we tested SV at 5dpf in *gucy1a1* mutants and found that in both, reduced SV persisted at later developmental stages (*Figure S7F-H, Movies S7,8*). To confirm that the observed phenotype reflects disruption of a shared Cygb2-NO-sGC signaling pathway in both *cygb2^801a^ and gucy1a1^9bp^* mutants, we tested whether pharmacological activation of sGC in *cygb^801a^* mutants could rescue cardiac function. Treatment of *cygb^801a^* mutants with the sGC activator Bay58 from 2ss to 34hpf fully restored SV (*Figure 4F-H),* confirming a critical role for sGC signaling in ventricular formation. Notably, neither DETA nor Bay58 rescued stroke volume in *gucy1a1^9bp^* mutants, as expected (*Figure S7C-D*). To assess a shared phenotype of increased ventricle wall thickness, measurements were performed on live embryos at 3dpf. Wall thickness at diastole and systole was increased in *gucy1a1^9bp^* (*Figure 4I-K, Movies S5,6*), similar to that observed in *cygb2^801a^* mutants (*Figure 4L-N, Movies S9,10*), indicating a conserved phenotype across mutant models and analytic approaches. To further determine whether NO-sGC signaling contributes to this phenotype, *cygb2^801a^* mutants were treated from 2ss to 34hpf with DETA and Bay58. These conditions fully restored ventricular wall thickness (*Figure 4L-N),* supporting a role for Cygb-NO-sGC in ventricular morphogenesis.

**Fig. 4.**
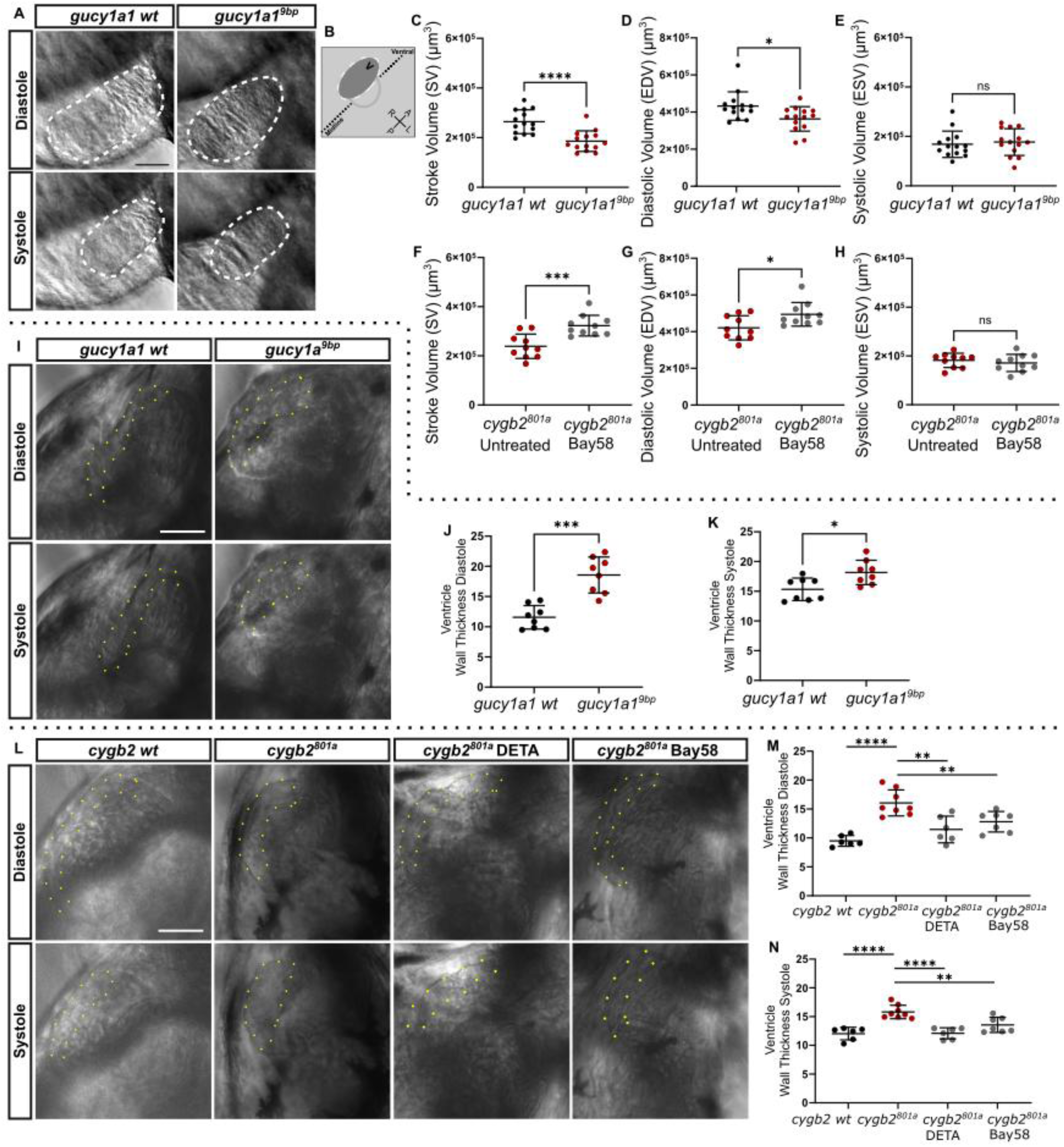
Ventricle area, ventricular diastolic volume, and stroke volume are reduced, and wall thickness is increased in sGC mutants. **(A)** Representative frames from DIC time-lapse imaging of live *gucy1a1wt* and *gucy1a1^9bp^* mutant embryos at 3 days post fertilization (dpf) showing volume at diastole and systole. Scale bar, 50µM. **(B)** Schematic of ventricle position and cardiac orientation. **(C-E)** Comparison of stroke volume, diastolic volume, and systolic volume between *gucy1a1wt*, and *gucy1a1^9bp^*. **(F-H)** Comparisons of stroke volume, diastolic volume, and systolic volume at 3dpf in live untreated and sGC activator (Bay58) treated *cygb2^801a^*;Tg(*myl7*:EGFP) embryos using fluorescent time-lapse imaging, treatments starting at 2 somite stage (ss). **(I,L)** Frames from live DIC time-lapse imaging of *gucy1a1wt*, *gucy1a1^9bp^*, *cygb2wt* and *cygb2^801a^* mutants at 3 dpf, depicting ventricle wall thickness at diastole and systole. Scale bar, 50µM. **(L)** NO donor (DETA) and Bay58 treatment starting at 2ss. **(J,K,M,N)** Comparison of wall thickness at diastole and systole. Sample size (n) corresponds to independent embryos represented by individual data points on plots. Student t-Test; *ns*, not significant; *p<0.05; **p<0.01;***p<0.001; ****p<0.0001.

### NO-sGC signaling regulates early cardiac progenitor organization

To determine whether Cygb2 and Gucy1a1 act through a shared molecular pathway during early cardiogenesis, we analyzed key markers of the ALPM at 10–12ss, when early cardiac and endothelial progenitors first emerge. In wildtype embryos, *drl* expression appeared as bilateral short, symmetrical stripes flanking the midline, marking anterior mesoderm that gives rise to cardiac and vascular lineages. In both *cygb2^801a^* and *gucy1a1^9bp^* embryos, *drl* domains were elongated, thinner, and extended posteriorly, indicating disrupted convergence and defective compaction of the ALPM (*Figure 5A*). Similarly, *kdrl*, which labels endothelial and endocardial progenitors, was expressed bilaterally in compact domains close to the midline in wildtype embryos, but it was elongated, more widely separated, and extended posteriorly in *cygb2^801a^* and *gucy1a1^9bp^* mutants (*Figure S8*). This finding is consistent with delayed or uncoordinated migration of endothelial/endocardial precursors. Despite these spatial changes, signal intensity was not diminished, suggesting that endothelial specification may be largely preserved, while progenitor patterning appears shifted from anterior (endothelial/endocardial) toward more posterior (hematopoietic) mesodermal domains. Expression of the early cardiac transcription factor *nkx2.5* was also altered at 10-12ss. In wildtype embryos, *nkx2.5* was expressed as two distinct, symmetrical domains within the ALPM, whereas in *cygb2^801a^* and *gucy1a1^9bp^* mutants, expression was frequently asymmetric or spatially disorganized (*Figure 5B*), suggesting loss of coordination between left and right cardiac fields. Importantly, wildtype siblings generated during establishment of the maternal-zygotic *cygb2^801a^* and *gucy1a1^9bp^* lines were outcrossed to minimize potential CRISPR off-target effects and displayed normal *drl*, *kdrl* and *nkx2.5* expression patterns at all developmental stages examined (*Figure S9*). The *drl* expression patterning abnormalities were rescued in *cygb2^801a^* embryos treated with either DETA or Bay58 (*Figure 5A*), confirming that early ALPM organization and synchronized cardiac specification depend on Cygb2-mediated NO–sGC signaling. Additionally, *nkx2.5* expression was also partially normalized by DETA treatment, although the strongest normalization was achieved in embryos treated with Bay58 (*Figure 5B*), confirming the importance of sGC signaling in this process. As *drl* marks early cardiac precursors fields and *nkx2.5* labels a broader population encompassing later myocardial progenitors, this differential rescue further supports the conclusion that NO–sGC signaling exerts its strongest influence during the earliest stages of cardiac precursor organization and initial heart structure formation.

**Fig. 5.**
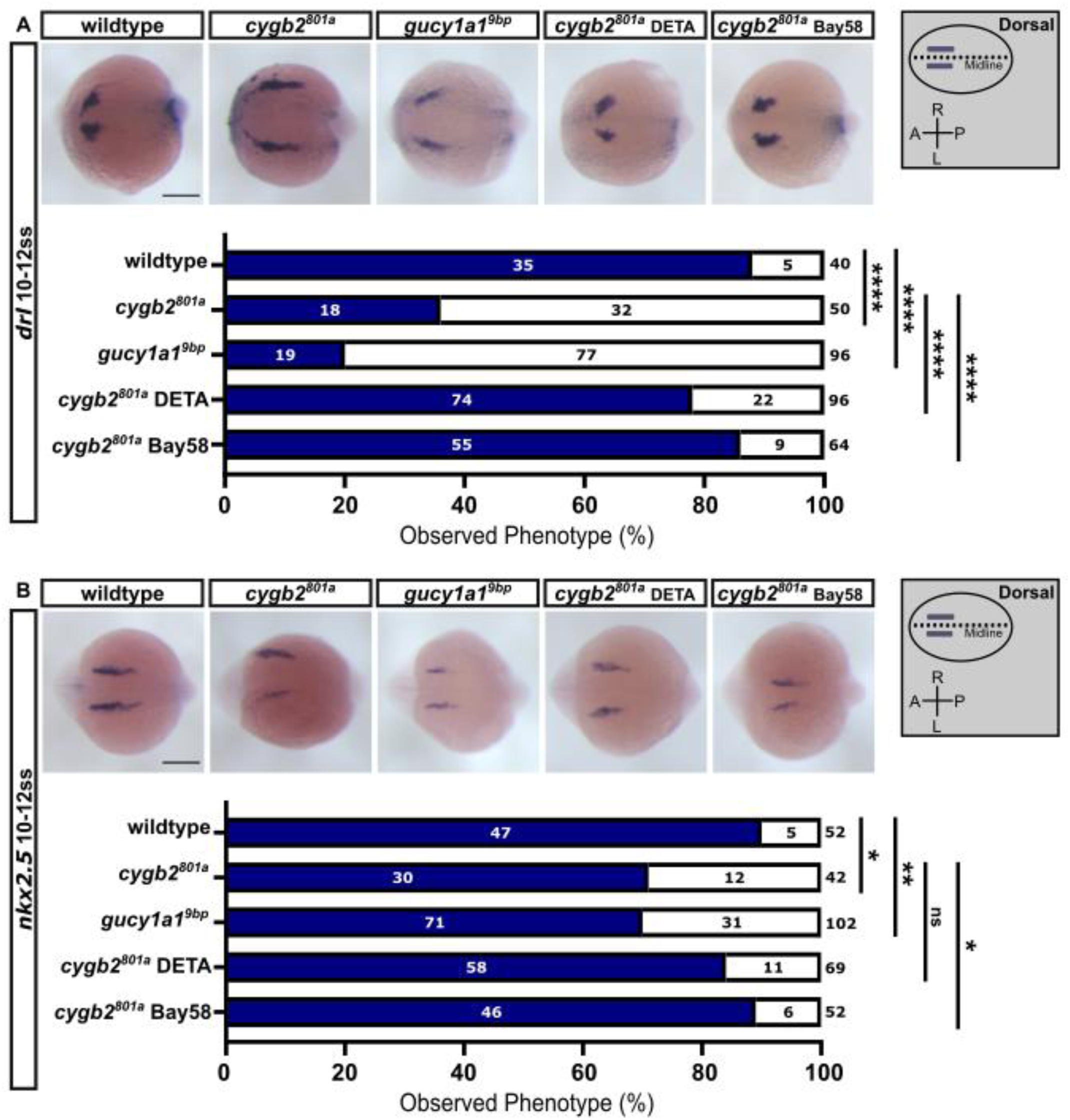
Early patterning of the anterior lateral plate mesoderm is disrupted in *cygb2* and *gucy1a1* mutants and rescued by NO–sGC activation. (A,B) Representative in situ hybridization images (top) and quantification of *drl* and *nkx2.5* expression patterns (bottom) in embryos at 10-12 somite stage. **(A)** *drl* normal expression pattern is quantified in solid-bars as percentage of embryos with distinct, short, and wide patches of bilateral *drl* expression at anterior edge of anterior lateral plate mesoderm (ALPM). In open-bars, percentage of embryos with elongated incongruous thin lines of bilateral *drl* expression along ALMP. **(B)** *nkx2.5* normal expression pattern is quantified in solid-bars as percentage of embryos with symmetrical, uniform, and distinct bilateral *nkx2.5* expression along ALMP. In open-bars, percentage of embryos with asymmetrical, patchy, or faint bilateral *nkx2.5* expression along ALMP. Number of embryos ranked is listed within and on top of bars. Treatments starting at shield. Scale bar, 200µm. On the right, schematic of ALPM anatomy with embryonic orientation. Fishers exact test, ns - not significant ≥ 0.05, * P≤ 0.05, ** P≤ 0.01, *** P≤ 0.001, **** P≤ 0.0001.

### NO–sGC signaling rescues ventricular gene expression

To assess whether early ALPM patterning defects persist during later stages of cardiogenesis, we examined *drl*, and *nkx2.5* expression at 28 hpf, when the heart tube has formed and endothelial and myocardial differentiation are underway. In wildtype embryos, *drl* expression is confined to the developing vasculature and the endocardium; in contrast, *cygb2^801a^* and *gucy1a1^9bp^* embryos exhibited reduced or absent *drl* expression in the heart tube (*Figure 6A*). This pattern is consistent with the shift observed at earlier stages from anterior (endothelial/endocardial) toward more posterior (hematopoietic) mesodermal domains. Consistently, *nkx2.5* expression in the ventricle was markedly reduced in both *cygb2^801a^* and *gucy1a1^9bp^* embryos, indicating impaired myocardial contribution to this region (*Figure 6B*). Treatment of *cygb2^801a^* embryos with DETA or Bay58 restored *drl* and *nkx2.5* expression patterns to near wildtype levels (*Figure 6A-B*). Together, these findings show that *cygb2* and *gucy1a1* mutants share a common set of early and late ALPM abnormalities that can be corrected by restoring NO-sGC signaling, linking early ALPM mis-patterning to the later emergence of a compact, hypoplastic ventricle.

**Fig. 6.**
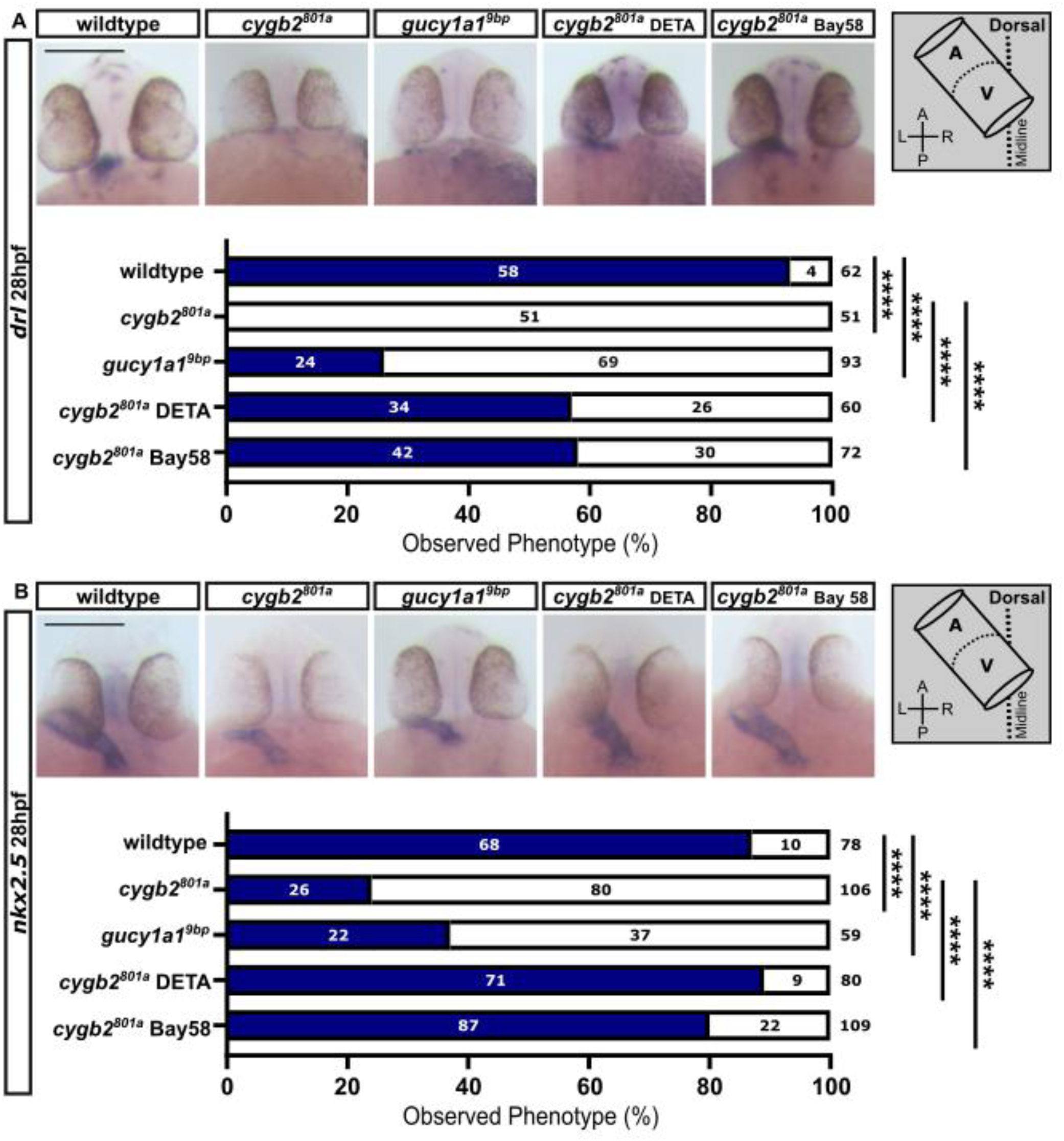
Disrupted specification of endothelial and myocardial lineages persists at 28 hours post fertilization and is normalized by restoring NO–sGC signaling. (A,B) Representative in situ hybridization images (top) and quantification of *drl* and *nkx2.5* expression patterns (bottom) in embryos at 28 hours post fertilization. (A) *drl* normal expression pattern is quantified in solid-bars as percentage of embryos with expression in the anterior region of the heart tube. In open-bars, percentage of embryos with complete loss of expression of *drl* in heart tube. (B) *nkx2.5* normal expression pattern is quantified in solid-bars as percentage of embryos with distinct clearly defined *nkx2.5* expression along the heart tube full-length. In open-bars, percentage of embryos with reduced or absent *nkx2.5* expression in a shortened heart tube. Number of embryos ranked is listed within and on top of bars. Treatments starting at shield. Scale bar, 200µm. On the right, schematic of the heart tube anatomy with embryonic orientation. Fishers exact test, ns - not significant ≥ 0.05, * P≤ 0.05, ** P≤ 0.01, *** P≤ 0.001, **** P≤ 0.0001.

## 4. Discussion

Nitric oxide (NO) signaling is a central regulator of cardiovascular physiology, with established roles in vascular tone, myocardial contractility, and hemodynamic homeostasis.^33–35^ In contrast, the contribution of NO signaling to early cardiac morphogenesis has been far less explored. Here, we show that Cygb2-dependent NO–sGC signaling is required for ventricular formation during zebrafish heart development. Loss of *cygb2* leads to a reduced SV associated with decreased ventricle volume, increased myocardial wall thickness, elevated cellular density, and impaired diastolic filling. These defects arise from abnormal migratory behavior of cardiac precursors, which display reduced migratory efficiency, excessive cell-spreading, and subsequent hypercompaction within the forming heart tube. Importantly, genetic loss of sGC (*gucy1a1*) fully phenocopies the *cygb2* mutant phenotype, and pharmacological activation of NO-sGC signaling in *cygb2* mutants rescues ventricular volume, cardiac progenitor migration, myocardial organization within the forming heart tube, and diastolic function. Together, these data identify NO–sGC signaling downstream of Cygb2 as a central pathway coordinating myocardial progenitor dynamics, ventricular morphogenesis, and functional maturation.

Cardiac progenitor migration is a fundamental driver of heart tube assembly and ventricular morphogenesis and depends on tightly regulated cell polarity, adhesion turnover, and cytoskeletal dynamics.^36–39^ Disruption of these processes can profoundly alter chamber size and myocardial architecture.^40,41^ In this context, our data indicate that reduced NO–sGC signaling places cardiac progenitors into a mechanically altered migratory state. NO-sGC-cGMP signaling is known to regulate actomyosin contractility, focal adhesion turnover, and cortical tension across multiple cellular contexts^42–44^ and cytoglobin-dependent control of NO bioavailability represents a key upstream determinant of this pathway^13,14^. Attenuation of NO–sGC signaling during development is therefore predicted to stabilize cell–substrate and cell–cell adhesions, resulting in excessive cell spreading, reduced migratory efficiency, and impaired cellular rearrangements at the tissue level. When such mechanically altered cells enter the confined environment of the forming heart tube, increased adhesion and cortical tension can bias morphogenesis toward tissue compaction and elevated cellular packing as demonstrated in epithelial and mesenchymal morphogenetic models.^45,46^ These properties provide a mechanistic basis for the increased ventricular cell density observed in *cygb2* mutants, together with the resulting thick-walled ventricle and impaired diastolic function.

We previously demonstrated that Cygb2 localizes to motile cilia and regulates cilia-dependent left–right patterning through NO–sGC signaling during early zebrafish development,^21^ establishing that Cygb2-dependent NO–sGC signaling operates during early developmental windows critical for laterality specification. The present study extends this role to ventricular formation while showing that ventricular defects do not arise directly from altered left–right axis specification (*Figure S2*). However, ventricular morphogenesis may still depend on cilia-associated signaling while remaining independent of final laterality outcomes. Here, we found that pharmacological restoration of NO–sGC signaling rescues ventricular morphology only when administered during early windows encompassing laterality determination and early somitogenesis, whereas later treatment fails to restore ventricular size and stroke volume, indicating that ventricular morphogenesis is temporally linked to early cilia-associated signaling events. During this developmental window, we found that *drl* and *kdrl* expression is spatially displaced and expanded within the ALPM, phenotypes consistent with defective cardiac progenitor migration rather than primary loss of cardiogenic identity^36,38,39,41^. Notably, *nkx2.5*, normally expressed symmetrically by early somitogenesis^47^ is found asymmetric in *cygb2* mutants, suggesting altered organization of subsets of cardiac progenitors. Recent single-cell studies indicate that *nkx2.5* marks transcriptionally distinct progenitor populations within the ALPM^29^ supporting the idea that the disrupted signaling perturbs progenitor composition rather than uniformly impairing cardiac fate. Early migratory defects likely lead secondarily to impaired differentiation or maintenance programs later, as reflected by the reduction of *nkx2.5* and *drl* expression within ventricular territories at 28 hpf.^48,49^ Importantly, dysregulation of NKX2-5 is associated with several forms of congenital heart disease in humans, including ventricular septal defects, tetralogy of Fallot, and HLHS, underscoring the relevance of these findings to human cardiac pathogenesis.^47,50,51^

Sequencing studies have identified enrichment of rare variants in cilia and ciliopathy-associated genes in subsets of HLHS patient cohorts,^4^ supporting a framework in which cilia-dependent pathways contribute to disease susceptibility in a subset of patients. Within this framework, zebrafish models exhibiting reduced ventricular size with compact myocardium have emerged as informative systems for studying cardiac morphogenesis and HLHS-like phenotypes.^9,49^ In our *cygb2* mutant model, HLHS-like ventricular hypoplasia originates from early events concomitant with cilia-dependent processes involved in left-right patterning. Our data indicate two possible mechanistic pathways underlying this phenotype. In one scenario, impaired NO-sGC signaling caused by loss of Cygb2 directly disrupts the coordination of cardiac progenitor migration, regional specification, and ventricle expansion, leading to reduced ventricular growth. In a second, distinct scenario, loss of Cygb2 perturbs cilia-dependent processes that regulate early cardiac morphogenesis and myocardial growth, beyond their established role in cardiac laterality, through downstream effects on progenitor deployment and tissue remodeling,^50^ independent of left–right axis determination. These mechanisms are not mutually exclusive and may interact in the multigenic context of human HLHS, where combined defects in NO signaling and cilia-associated pathways could converge on shared morphogenetic processes controlling ventricular development. The *cygb2 and gucy1a1* models characterized here converge on the hallmark ventricular phenotype of HLHS while revealing a distinct initiating axis centered on Cygb-dependent NO–sGC signaling. Together with prior work demonstrating that modulation of NO bioavailability influences myocardial plasticity and regeneration,^52^ these findings establish NO–sGC signaling as a tractable developmental pathway whose disruption leads to ventricular hypoplasia and whose restoration is sufficient to rescue ventricular structure and function. Our data indicate that pharmacological activation of sGC, previously unexplored in the context of hypoplastic ventricular disease, can restore ventricular structure and function in HLHS-like phenotypes in vivo. These results position zebrafish as a valuable model for interrogating early developmental mechanisms underlying ventricular hypoplasia and for assessing sGC-targeted interventions in CHD.

## Author Contributions

Adam Clark, Paola Corti designed and conducted experiments, analyzed data, and wrote the manuscript. Paola Corti directed the research and conceptualization of experiments. Rasmus Hejlesen and Muddassar Iqbal reviewed the manuscript. Rasmus Hejlesen constructed a data set for transcript variance analysis that guided hypothesis generation. Weng Tzu-Ting conducted in situ hybridization experiments. Muddassar Iqbal designed, conducted; and analyzed data for siRNA knockdown, wound healing, and Western Blot experiments. Akueba Bruce conducted zebrafish husbandry and genotyping.

## Acknowledgements

We are thankful to Dr. Mark T. Gladwin (Dean, University of Maryland School of Medicine) for expert mentorship and guidance in project design. We thank Dr. Joseph Mauban, Dr. Shilpa Kumar and the University of Maryland School of Medicine Center for Innovative Biomedical Resources, Confocal Microscopy Core for invaluable assistance in imaging, and the Translational Genomics Lab for Sanger Sequencing services.

## Data availability

The data used to produce this article will be shared at reasonable request to the corresponding author.

