## Supplement for "Cytoglobin-dependent NO-sGC-cGMP signaling regulates ventricular morphogenesis and diastolic function"

Corti, Paola, PhD

Department of Biochemistry and Molecular Biology, University of Maryland School of Medicine, Baltimore, MD 21201. USA

**Supplementary Material**

Supplementary Tables page 2-3

Supplementary Figures page 4-14

Supplementary Movies page 15-16

### Supplementary Tables

| Primer Name | F/R | Sequence (5' – 3') | Assay |
|---|---|---|---|
| *Gucy1a1* gRNAx34 | F | ATTTAGGTGACACTATAGGCTCACACCAAGCCCCATGGTTTTAGAGCTAGAAATAGC | gRNA PCR |
| *Gucy1a1* gRNAx87 | F | ATTTAGGTGACACTATAGGACGGTCATATTCTCAGAGGTTTTAGAGCTAGAAATAGC | gRNA PCR |
| gRNA universal primer | R | AAAAGCACCGACTCGGTGCCACTTTTTCAAGTTGATAACGGACTAGCCTTATTTTAACTTGCTATTTCTAGCTCTAAAAC | gRNA PCR |
| *Gucy1a1* gucygRNAx34 | F | CCCCTGAGATACAGCGTTACAT | Genotyping |
| *Gucy1a1* gucygRNAx34 | R | TATCCTGGAAGCTGGCTCGTA | Genotyping |
| *Gucy1a1* gucygRNAx87 | F | CCCCGTAGCATGCAGCTCTGA | Genotyping |
| *Gucy1a1* gucygRNAx87 | R | CTCAGGGGTGGGTGGTGTGC | Genotyping |
| *Cygb2*_Ex1 | F | CACAGCCTCTCATTTCACCA | Genotyping |
| *Cygb2*_Ex1 | R | ATGATGGATAAATTTGCACACTG | Genotyping |
| *drl-rProbe* | F | GATTCTCACATAAAGCGCACC | ISH |

| | | | |
|---|---|---|---|
| *drl -rProbe* | R | TAATACGACTCACTATAGGGAGACAGAGTGAACACATGTAAGGTCT | ISH |
| *kdrl -rProbe* | F | AGCTGCCCAATGTTTCCAGA | ISH |
| *kdrl -rProbe* | R | TAATACGACTCACTATAGGGAGATCTTCCTCCGAGTCGCAGTA | ISH |
| *nkx2.5 -rProbe* | | Plasmid; Generous gift from the Beth Roman Lab | ISH |
| *gucy1a1* 1000 | F | CGAGCCAGCTTCCAGGATAT | ISH |
| *gucy1a1* 1000 | R | TAATACGACTCACTATAGGGAGATCTTTTGATCGTGAGCCTGGA | ISH |
| *gucy1a1* 300 | F | TCTCGGGTATCATCCAGGCG | ISH |
| *gucy1a1* 300 | R | TAATACGACTCACTATAGGGAGACTCTGCATCTCCCTAGCCGT | ISH |
| *si-CYGB 13.1* | | GUCCUCCUUUCUAGUUU | siRNA |
| *si-CYGB 13.2* | | AUAAACCAGCCCCGUUUC | siRNA |

**Supplemental Table 1. Primers, oligos, and silencing RNAs used in this study**

**Supplementary Figures**

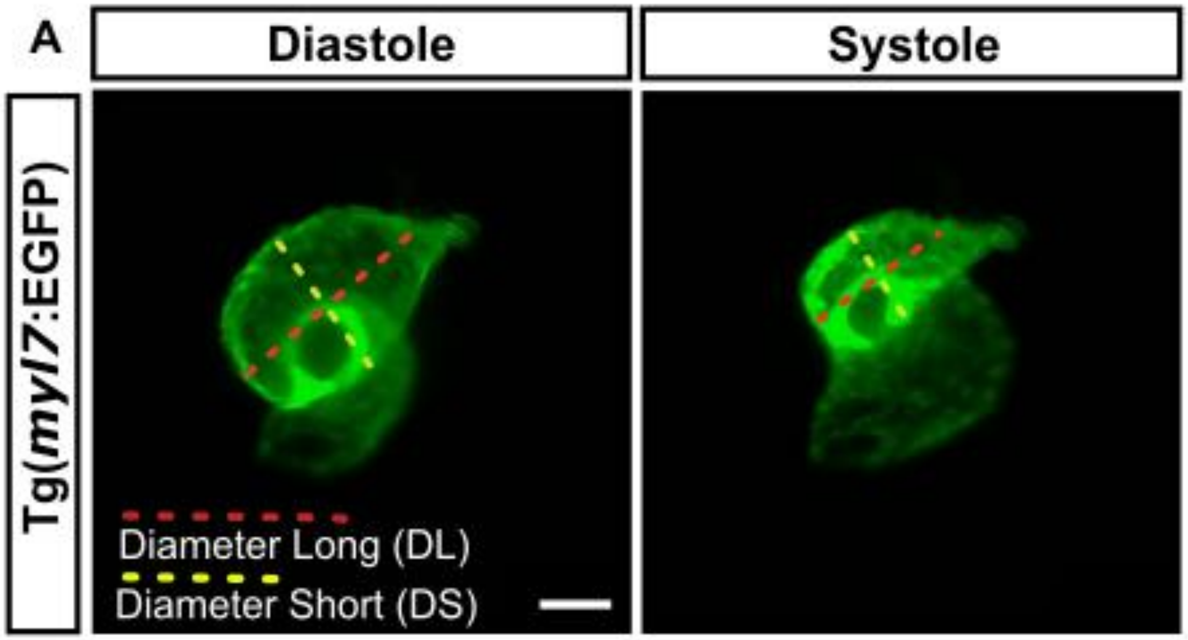


**Supplemental Figure 1. Analysis of stroke volume using the prolate spheroid method. (A)** Representative frames from fluorescent time-lapse imaging of live wild type embryos in the Tg(*myl7*:EGFP) transgenic background (EGFP, green) at 3 days post fertilization showing references to measurements of diameter long and short used in ventricle volume calculations. Scale bar, 50μM.

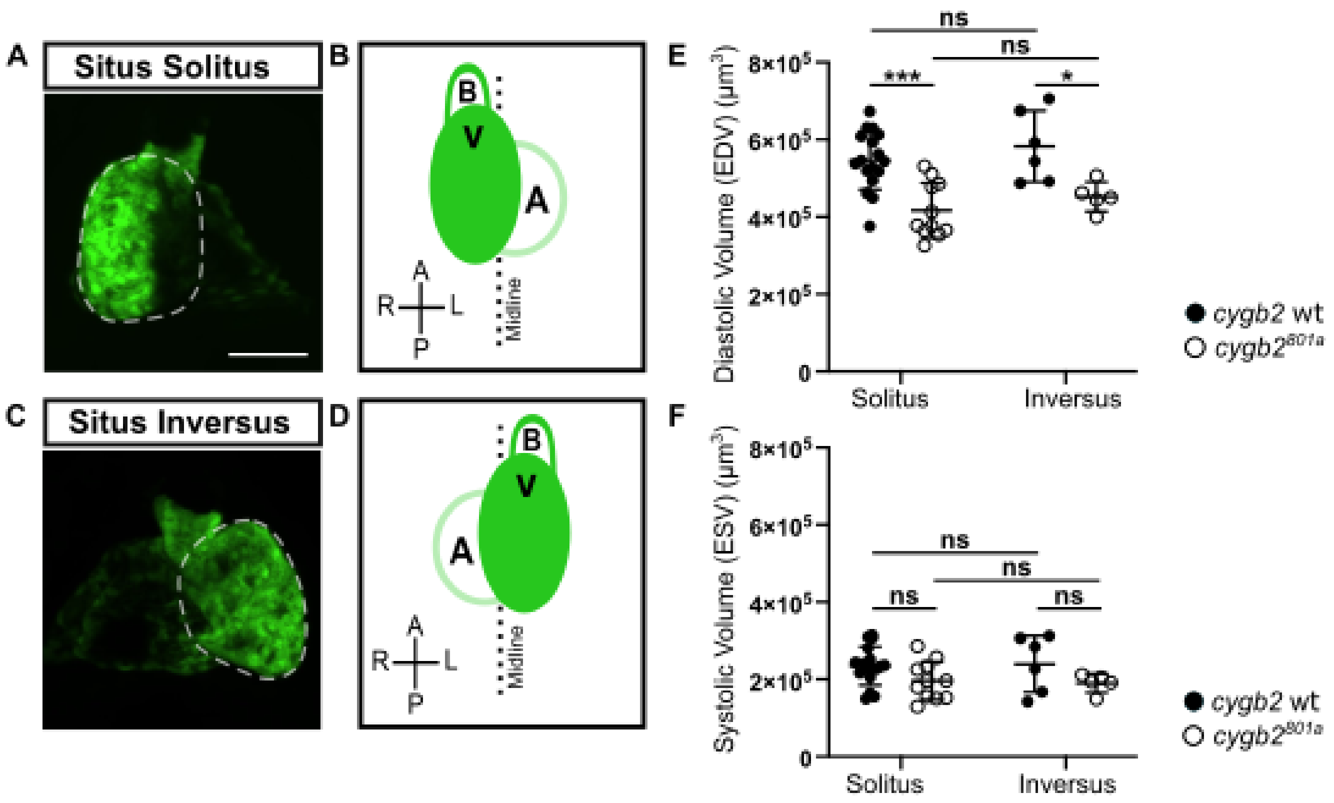


**Supplemental Figure 2. Cardiac laterality does not effect stroke volume. (A,C)** Representative whole-mount confocal projections of *cygb2*$^{801a}$;Tg(*myl7*:EGFP) embryos at 3 days post fertilization (EGFP, green) showing normal and reversed ventricles (*situs solitus* and *situs inversus)*. Scale bar, 50µm. **(B,D)** Schematics illustrating chamber anatomy and cardiac laterality. A-atrium; V-ventricle; B-bulbus arteriosus **(E,F)** Effect of *situs solitus* and *situs inversus* on stroke volume. Sample size (n) corresponds to independent embryos represented by individual data points on plots. Students t-Test; ns, not significant, * P≤ 0.05, ** P≤ 0.01, *** P≤ 0.001.

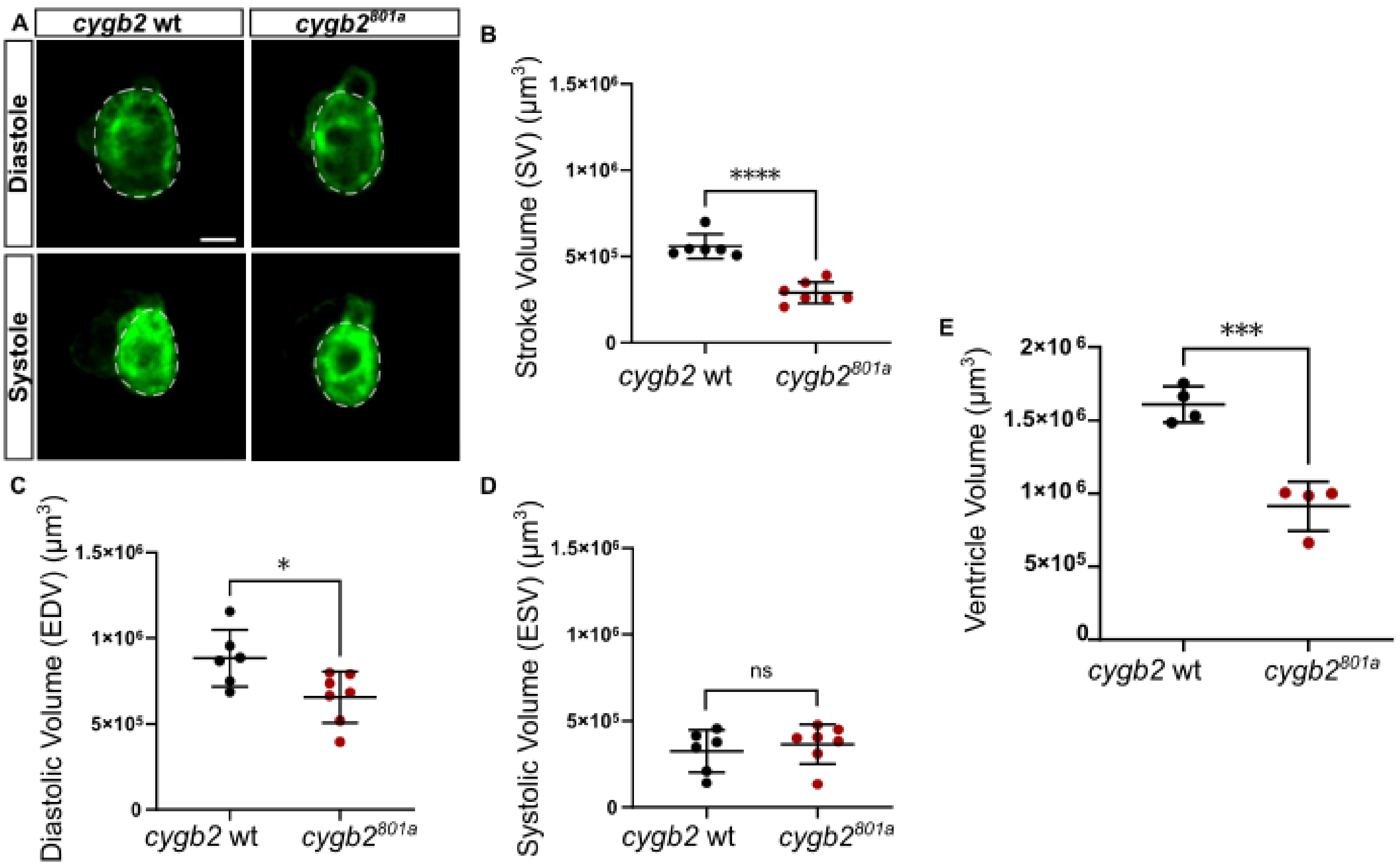


**Supplemental Figure 3. Decreased stroke volume and reduced ventricle size persist at 5 days post fertilization in *cygb2* mutants. (A)** Representative frames from fluorescence time-lapse imaging of live *cygb2*wt*;*Tg(*myl7*:EGFP) and $cygb2^{801a}$;Tg(*myl7*:EGFP) embryos at 5 days post fertilization. Scale bar, 50µm. **(B-D)** Comparisons of stroke volume at 5 days post fertilization. **(E)** Comparisons of ventricle volume in whole-mount *cygb2*wt*;*Tg(*myl7*:EGFP) and $cygb2^{801a}$;Tg(*myl7*:EGFP) measured by Lightsheet confocal imaging. Sample size (n) corresponds to independent embryos represented by individual data points on plots. Student t-Test; ns, not significant, * P≤ 0.05, *** P≤ 0.001, **** P≤ 0.0001.

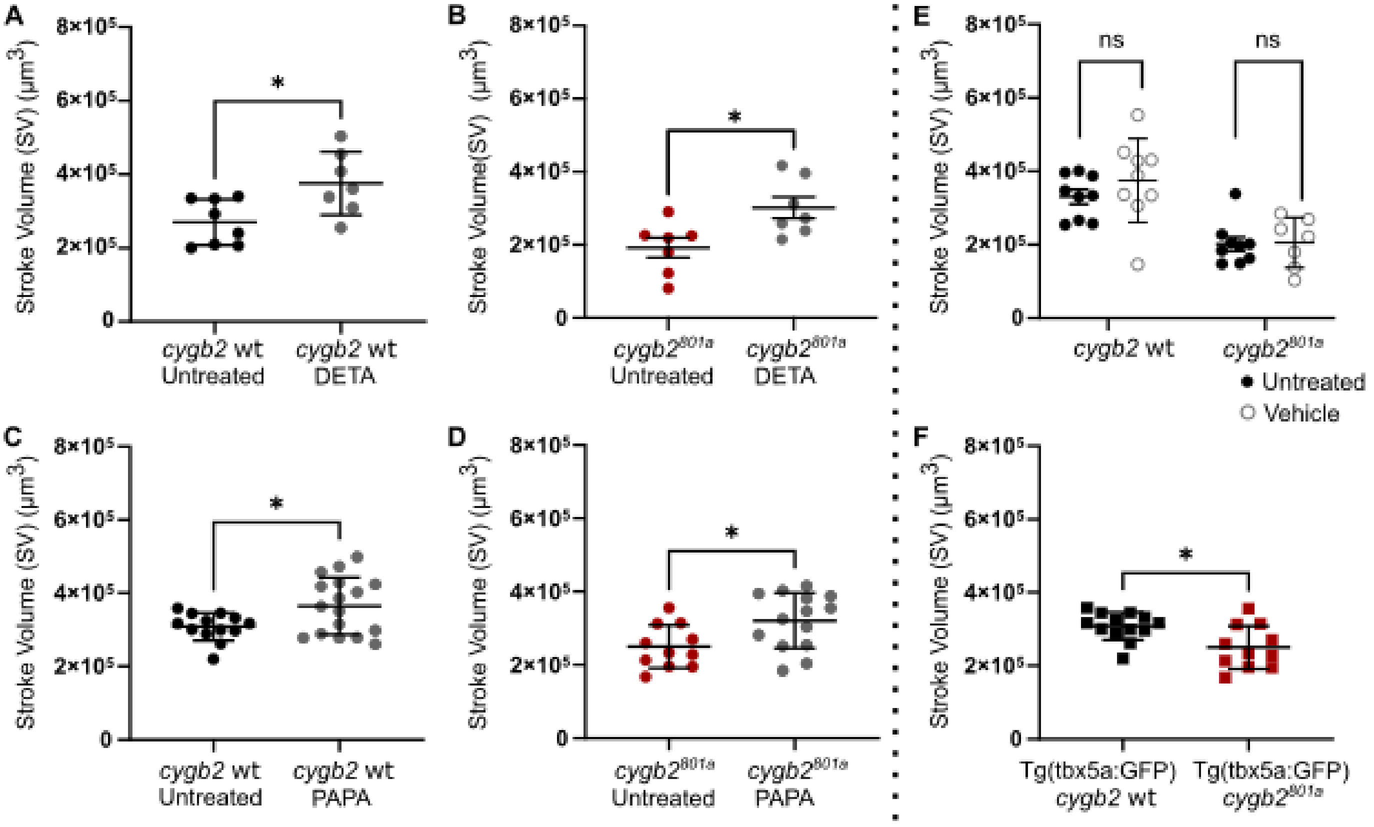


**Supplemental Figure 4. Acute treatment with NO donors DETA NONOate and PAPA NONOate increases stroke volume in *cygb2* wildtype and *cygb2* mutant. (A-E)** Stroke volume (SV) comparisons between live *cygb2*wt;Tg(*myl7*:EGFP) and $cygb2^{801a}$;Tg(*myl7*:EGFP) at 3 days post fertilization (dpf), measured from fluorescent time-lapse imaging. (**A-D)** Comparison of SV between untreated and NO donor-treated (DETA, PAPA) embryos, 10-minute treatment starting at 78 hours post fertilization. **(E)** Effect of vehicle control sodium hydroxide on SV. **(F)** Comparisons of SV between *cygb2wt;*Tg(*tbx5a*:EGFP);Tg(*tcf21*:dsRED) and $cygb2^{801a}$;Tg(*tbx5a*:EGFP);Tg(*tcf21*:dsRED) at 3dpf measured using live fluorescence time-lapse imaging. Sample size (n) corresponds to independent embryos represented by individual data points on plots. Student t-Test; ns, not significant, * P≤ 0.05.

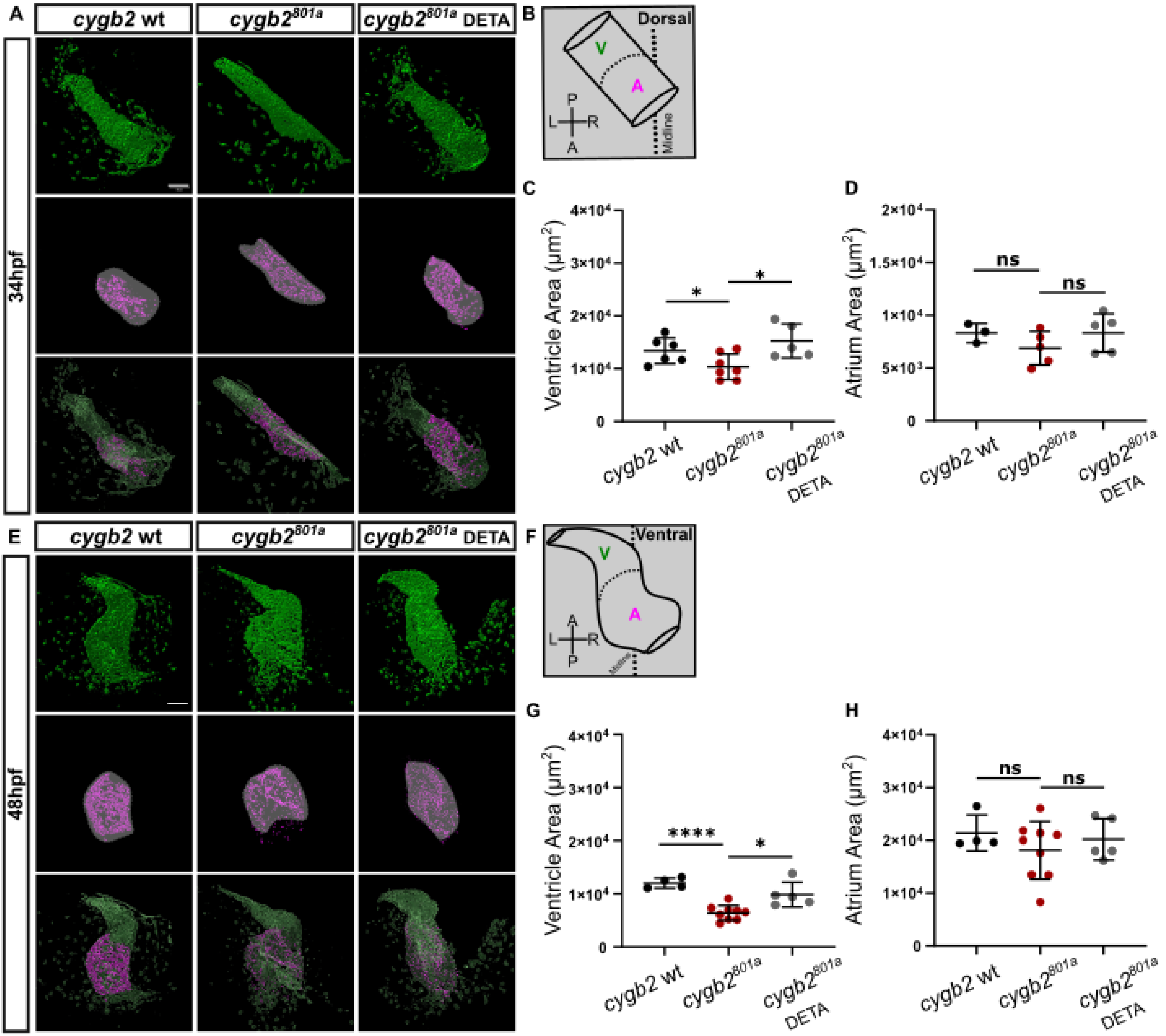


**Supplemental Figure 5. Ventricular domain is reduced in *cygb2* mutant at 34-48 hours post fertilization. (A,E)** Representative 3D surface renderings from confocal z-series of *cygb2wt*;Tg(*myl7*:EGFP) and *cygb2$^{801a}$*;Tg(*myl7*:EGFP) heart tube at 34 and 48 hours post fertilization, immune-labeled for Myh6 (GFP, green; Myh6, magenta). Scale bar, 50μM, treatment starting at 2 somite stage. **(B,F)** Schematic illustrating heart tube anatomy. A-atrium; V-ventricle (**C,D,G,H**) Comparisons of ventricular and atrial domains. Sample size (n) corresponds to independent embryos represented by individual data points on plots. Student t-Test; ns, not significant, * P≤ 0.05, ** P≤ 0.01, *** P≤ 0.001, **** P≤ 0.0001.

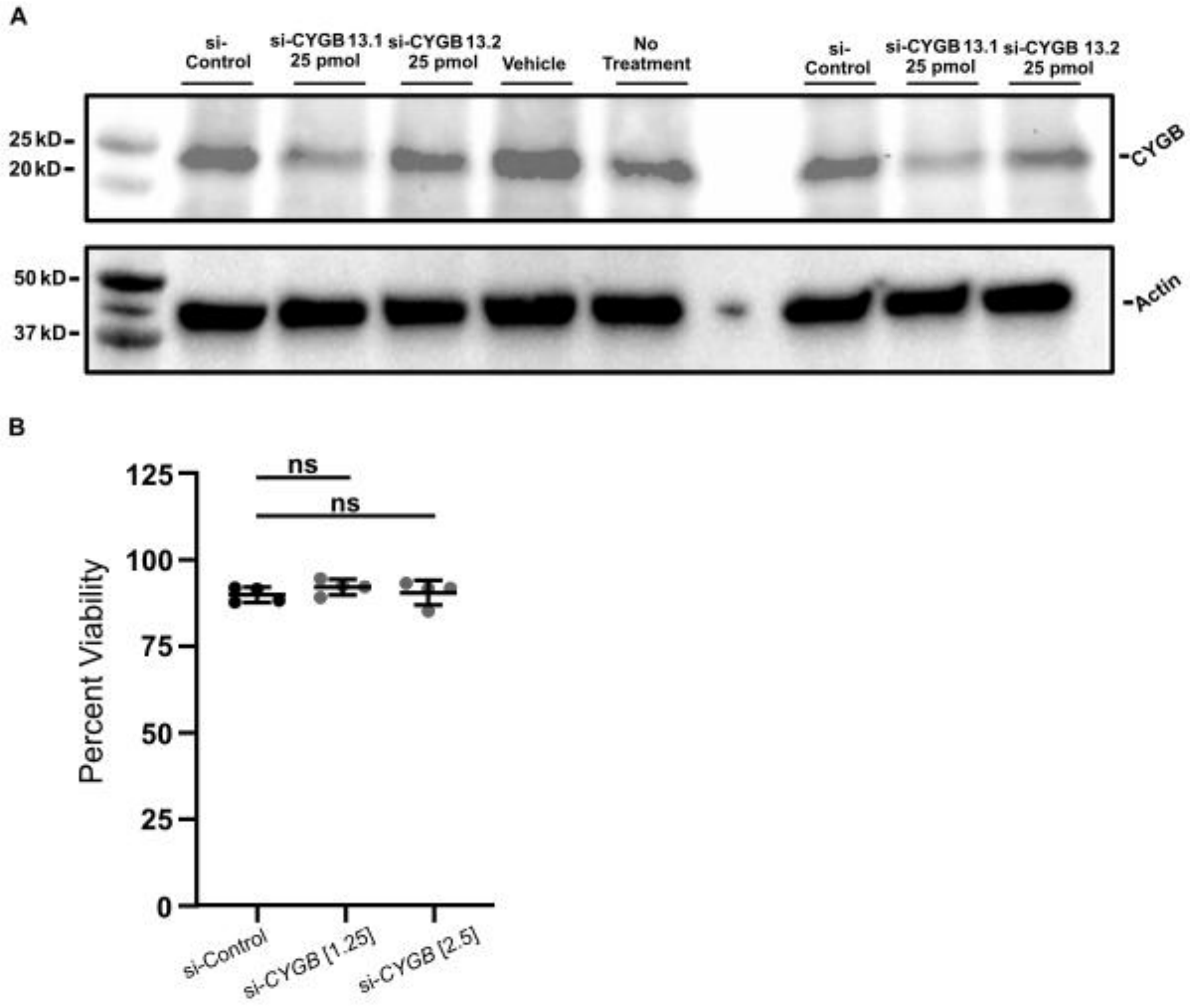


**Supplemental Figure 6. Cytoglobin protein expression in A549 cultures is reduced using small interfering RNA. (A)** Representative images of Western Blots bands depicting CYGB protein expression in A549 cells transfected with non-targeting siRNA, human *CYGB* targeting siRNAs (si-CYGB 13.1 and si-CYGB 13.2), lipofectamine (vehicle), and non-transfected (no treatment), along with β-Actin housekeeping control. **(B)** Comparison of cell viability in non-targeting and si-CYGB 13.1 at

1.25 pmol and 2.5 pmol concentrations. Sample size (n) corresponds to independent cultures represented by individual data points on plot. Student t-Test; ns, not significant,

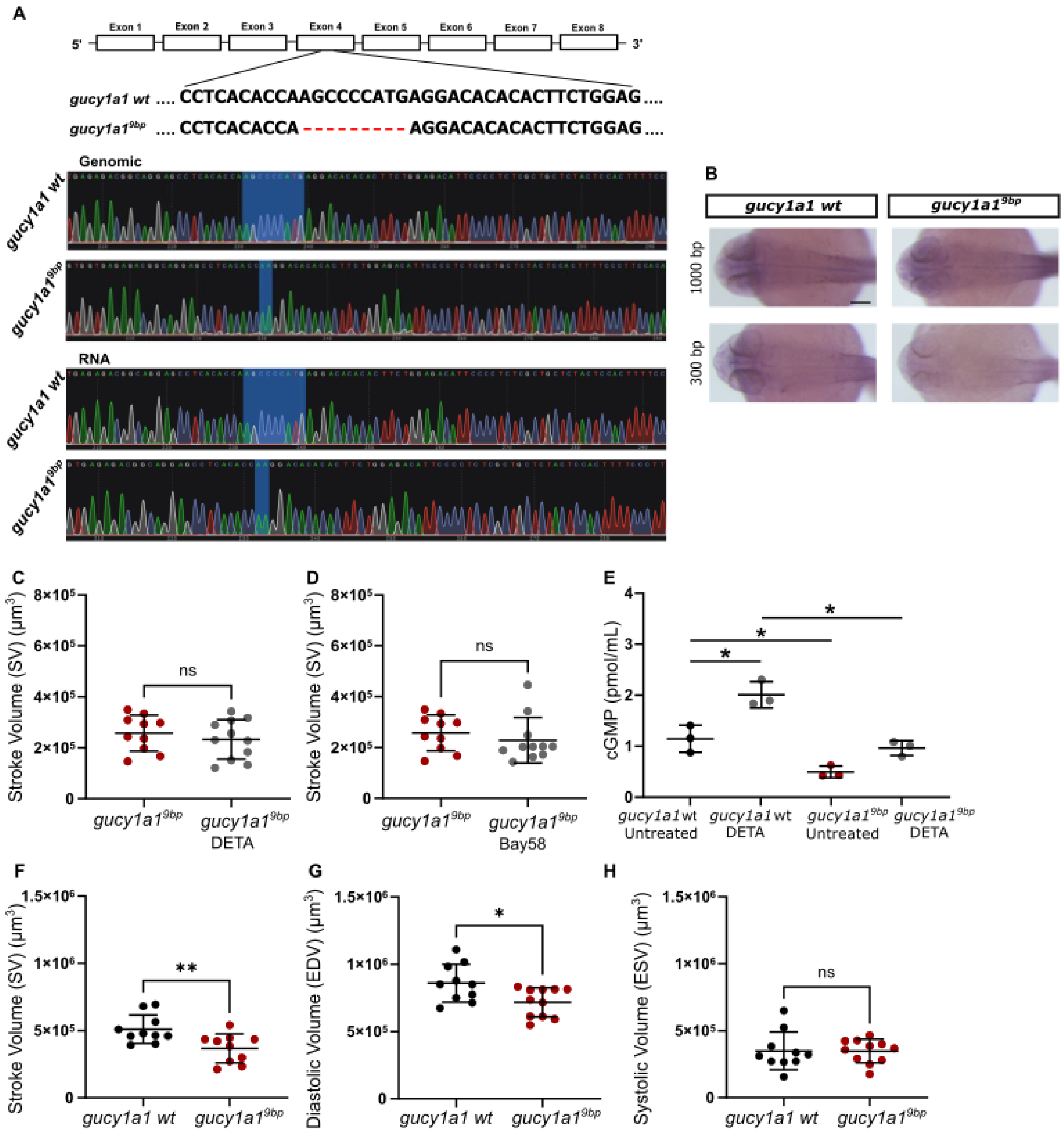

**Supplemental Figure 7. Generation and functional characterization of *gucy1a1* mutants. (A)** Schematic illustrating CRISPR/Cas9 mediated genome editing of *gucy1a1* resulting in a 9 base pair deletion in exon 4 (Top), chromatograms from Sanger sequencing of genomic DNA and cDNA of *gucy1a1*wt and $gucy1a1^{9bp}$ (Bottom). Shaded region in *gucy1a1*wt depicts the 9 nucleotide sequence missing in $gucy1a1^{9bp}$. The shaded region in $gucy1a1^{9bp}$ depicts alanine nucleotides flanking up and downstream of 9 base pair deletion. **(B)** Representative in situ hybridization images using 1000 and 300 base pair *gucy1a1* riboprobes. Scale bars, 200µm. Sample size (n) corresponds to independent embryos imaged per condition, n = 6. **(C-D)** Comparison of stroke volume (SV) between *gucy1a1*wt, $gucy1a1^{9bp}$, $gucy1a1^{9bp}$ DETA and Bay58 treated embryos at 3 days post fertilization (dpf). Treatments starting at 2 somite stage. **(E)** Comparison of cGMP levels in untreated and NO donor (DETA) treated *gucy1a1*wt and $gucy1a1^{9bp}$ at 28 hours post fertilization. **(F-H)** Comparisons of SV in live *gucy1a1*wt and $gucy1a1^{9bp}$ embryos at 5dpf measured using DIC time-lapse imaging. Sample size (n) corresponds to independent embryos represented by individual data points on plots. Student t-Test; ns, not significant, * P≤ 0.05, ** P≤ 0.01.

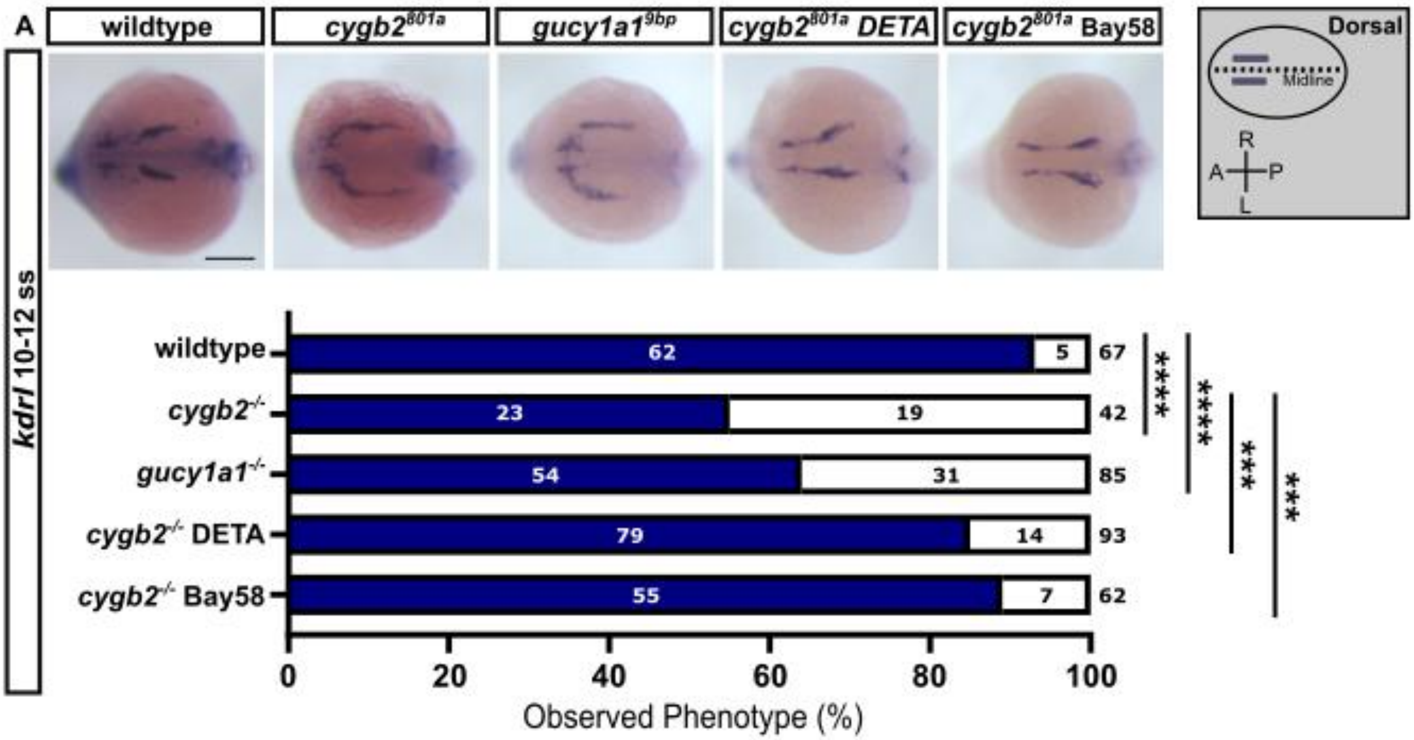


**Supplemental Figure 8. Disrupted patterning of *kdrl* in the anterior lateral plate mesoderm in *cygb2* and *gucy1a1* mutants is rescued by NO–sGC activation in *cygb2* mutants. (A)** Representative in situ hybridization images of wildtype, *cygb2*$^{801a}$, *gucy1a1*$^{9bp}$, *cygb2*$^{801a}$ NO donor (DETA), and sGC activator (Bay58) treated embryos at 10-12 somite stage. *kdrl* normal expression pattern is quantified in solid-bars as percent of embryos with bilateral *kdrl* expression forming a V-shaped point at anterior edge of anterior plate mesoderm (ALMP). In open-bars, percent of embryos with U-shaped point at anterior edge of the ALMP. Number of embryos ranked is listed within and on top of bars. Treatments initiated at shield. Scale bar, 200µm. On the right, schematic of ALPM anatomy with embryonic orientation. Fishers exact test, *** P≤ 0.001, **** P≤ 0.0001.

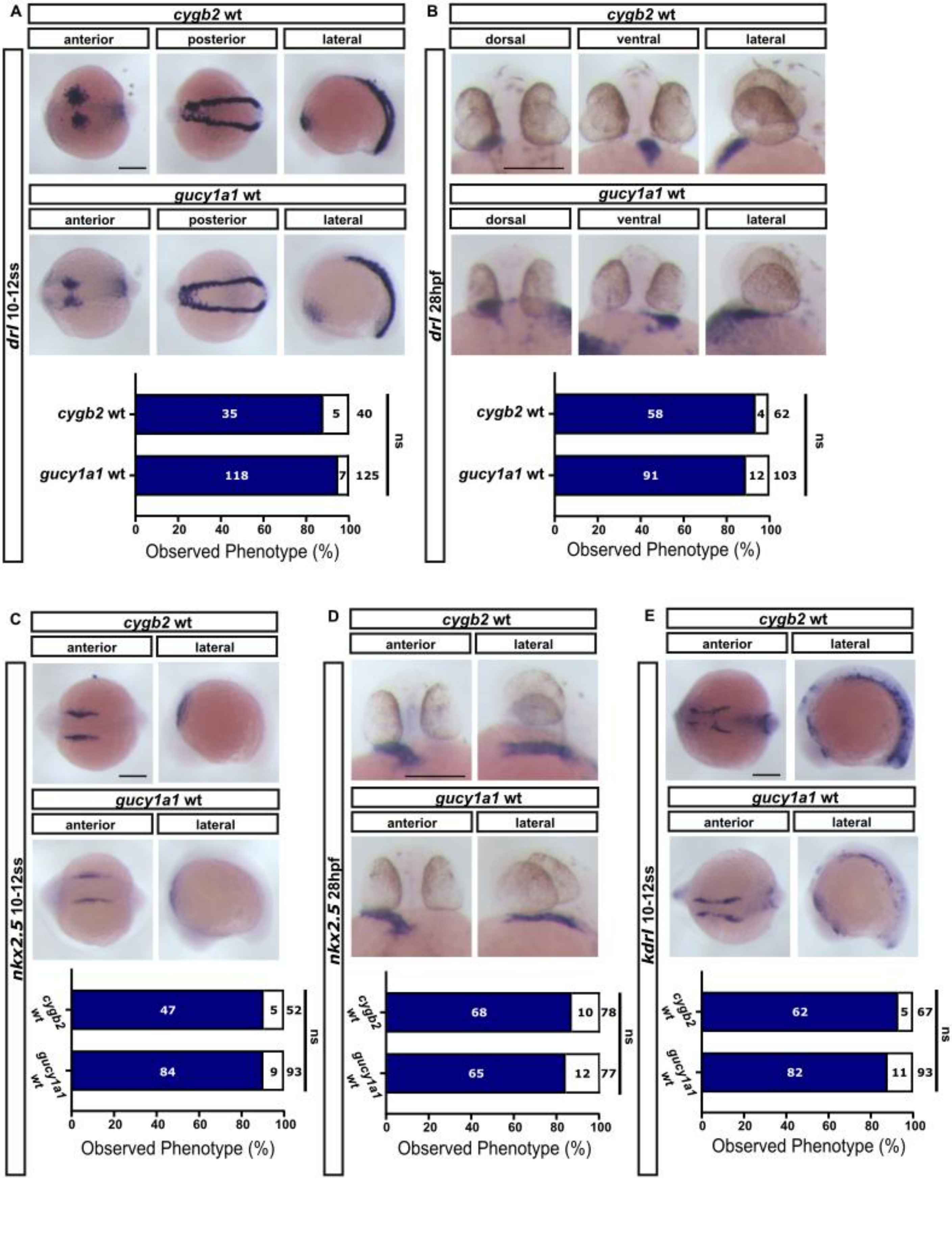

A
cygb2 wt
anterior
posterior
lateral
gucy1a1 wt
drl 10-12ss
35
5
40
118
7
125
ns
0 20 40 60 80 100
Observed Phenotype (%)
B
dorsal
ventral
drl 28hpf
58
4
62
91
12
103
C
nkx2.5 10-12ss
47
5
52
84
9
93
D
nkx2.5 28hpf
68
10
78
65
12
77
E
kdrl 10-12ss
62
5
67
82
11
93


151

**Supplemental Figure 9. Early patterning of the anterior lateral plate mesoderm and elongated heart tube is consistent between *cygb2* and *gucy1a1* wildtype siblings. (A-E)** Representative in situ hybridization images of *cygb2*wt, and *gucy1a1wt* embryos at 10-12 somite stage and 28 hours post fertilization showing expression patterning of *drl, nkx2.5, and kdrl*. In bar charts, normal expression pattern is shown in solid-bars as percent of total; open-bars indicates the percentage of embryos with any expression patterning deviating from the wildtype. The number of embryos ranked is listed within and on top of bars. Scale bar 200μm. Fishers exact test, ns - not significant ≥ 0.05.

**Supplementary Movies**

Movies (1-10) - Movie speeds normalized to 10 frames per second, scale bars 50μm.

Movies (1-4) - Representative fluorescence time-lapse videos of cardiac cycle phases.

Movies (5-10) - Representative DIC time-lapse showing cardiac cycle phases and ventricle wall thickness.

File Name: Supplementary Movie 1

Description: 3dpf *cygb2wt*;Tg(*myl7*:EGFP) embryos.

File Name: Supplementary Movie 2

Description: 3dpf *cygb2*$^{801a}$;Tg(*myl7*:EGFP) embryos.

File Name: Supplementary Movie 3

Description: 5dpf *cygb2wt*:Tg(*myl7*:EGFP) embryos.

File Name: Supplementary Movie 4

Description: 5dpf *cygb2*$^{801a}$:Tg(*myl7*:EGFP) embryos.

File Name: Supplementary Movie 5

Description: 3dpf *gucy1a1*wt embryos.

File Name: Supplementary Movie 6

Description: 3dpf *gucy1a1*$^{9bp}$ embryos.

File Name: Supplementary Movie 7

Description: 5dpf *gucy1a1*wt embryos.

File Name: Supplementary Movie 8

Description: 5dpf *gucy1a1*$^{9bp}$ embryos.

File Name: Supplementary Movie 9

Description: 3dpf *cygb2*wt embryos.

File Name: Supplementary Movie 10

Description: 3dpf *cygb2*$^{801a}$ embryos.
